# Dissociation of tonotopy and pitch in human auditory cortex

**DOI:** 10.1101/2020.09.18.303651

**Authors:** Emily J. Allen, Juraj Mesik, Kendrick N. Kay, Andrew J. Oxenham

## Abstract

Frequency-to-place mapping, or tonotopy, is a fundamental organizing principle from the earliest stages of auditory processing in the cochlea to subcortical and cortical regions. Although cortical maps are referred to as tonotopic, previous studies employed sounds that covary in spectral content and higher-level perceptual features such as pitch, making it unclear whether these maps are inherited from cochlear organization and are indeed tonotopic, or instead reflect transformations based on higher-level features. We used high-resolution fMRI to measure BOLD responses in 10 participants as they listened to pure tones that varied in frequency or complex tones that independently varied in either spectral content or fundamental frequency (pitch). We show that auditory cortical gradients are in fact a mixture of maps organized both by spectral content and pitch. Consistent with hierarchical organization, primary regions were tuned predominantly to spectral content, whereas higher-level pitch tuning was observed bilaterally in surrounding non-primary regions.

## INTRODUCTION

A key organizing principle of the auditory system is tonotopy, an orderly mapping of sound frequency to place. Tonotopy is established in the cochlea, where different frequencies maximally displace different locations along the basilar membrane, in a high-to-low ordering from the base to the apex (Von Békésy, 1960). This tonotopic organization has been found at numerous stages of the auditory pathways up to and including auditory cortex (e.g., Von Békésy, 1960; Moerel et al., 2015; Saenz and Langers, 2014). Studies of cortical mapping using fMRI have typically employed pure tones or narrowband noises (e.g., Da Costa et al., 2011; Formisano et al., 2003; Saenz and Langers, 2014; Striem-Amit et al., 2011; Talavage et al., 2004) in much the same way as has historically been done to establish tonotopy in earlier stages of the auditory processing hierarchy (Von Békésy, 1960; Bourk et al., 1981; Cooper, 1999; Narayan et al., 1998; Nuttall and Dolan, 1996; Ruggero et al., 1997; Schreiner et al., 1997). However, pure tones, narrowband stimuli, and even many natural sounds, conflate two primary perceptual attributes of sound: pitch height and timbral brightness. In most sounds, pitch is determined by the fundamental frequency (F0), whereas the sound quality, or timbre, is affected by the spectral centroid (Fc) of the sound’s energy distribution (Allen and Oxenham, 2014; Krumhansl and Iverson, 1992; Marozeau et al., 2003). Because previous studies have used stimuli in which pitch height and timbral brightness covary, it remains unclear whether the spatial organization observed in cortex actually reflects frequency-to-place mapping, inherited from the cochlear representation of spectral content, or whether some or all portions of the cortical maps instead reflect one or more higher-level features.

There is mounting evidence for hierarchical organization within auditory cortex, with primary areas near Heschl’s gyrus showing a preference for relatively simple acoustic features, and surrounding non-primary areas showing greater sensitivity to complex auditory objects, such as speech and music (de Heer et al., 2017; Kell et al., 2018; Norman-Haignere et al., 2015). Thus, it may be that the multiple gradients identified in previous studies as multiple tonotopic maps (e.g., Moerel et al., 2014; Saenz and Langers, 2014), may in fact reflect different maps of different auditory features.

To distinguish the mapping of spectral variation and pitch in cortical representations, we used high-resolution functional magnetic resonance imaging (fMRI) to measure cortical responses to sound sequences that varied systematically and orthogonally in F0 and Fc and used computational models to characterize the spatial organization of each of these features. Our results demonstrate that the well-documented cortical tonotopy is primarily driven by spectral content, consistent with the organization found in the more peripheral auditory pathways. However, they also reveal bilateral maps of pitch that partially overlap with the tonotopic maps but are located primarily outside Heschl’s gyrus. Overall, the findings reveal the existence of multiple spatially organized maps that reflect both tonotopy and higher-level pitch bilaterally within human auditory cortex.

## RESULTS

### Behavioral Results

While undergoing fMRI, participants (n = 10) listened to sequences of pure tones that varied in frequency as well as sequences of complex tones that varied in either Fc or F0, referred to as the pure-tone, timbre, and pitch conditions, respectively (Figure 1). To ensure that the participants remained alert and attentive, they were instructed to indicate via button box whether the current tone was higher or lower than the previous one. Behavioral performance was high across all three conditions for all participants, suggesting that they successfully attended to the stimuli. The average percentage of correct responses was 96.8% [SD = 2.3%] in the pure-tone condition, 93.1% [4.5%] in the timbre condition, and 95.8% [4.4%] in the pitch condition. Due to near-ceiling performance in all conditions, a non-parametric Friedman test was run to detect differences in performance between conditions, which revealed a significant main effect χ^2^(2) = 9.6, *p* = 0.008. Post-hoc analysis with a Wilcoxon signed rank test was then conducted to compare conditions. After a Bonferroni correction for multiple comparisons, setting α to 0.017 (0.05/3), no significant differences were found between any of the conditions (pitch vs. timbre: *Z* = −1.78, *p* = 0.074; pitch vs. pure tones: Z = 0.00, *p* = 1.00; timbre vs. pure tones: *Z* = −2.35, *p* = 0.019). Therefore, differences in cortical representations between the three conditions are unlikely to be due to differences in behavioral performance.

**Figure 1.**
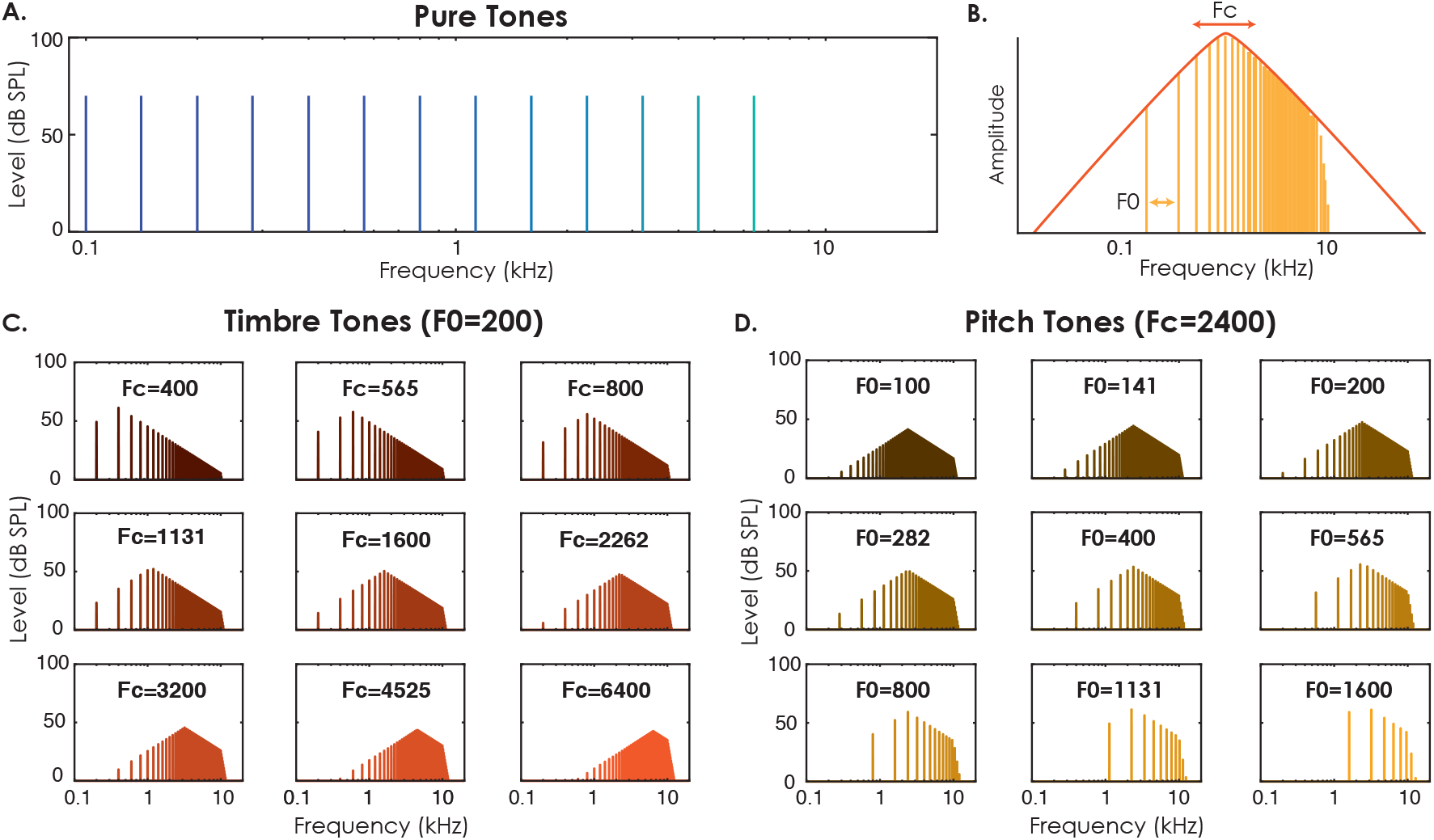
Stimulus Configurations. (A) Frequencies of all thirteen tones used in the pure-tone condition. (B) Schematic diagram of harmonic complex tone manipulation. Shifts in the spectral envelope to the right or left (orange) correspond to increases or decreases in spectral centroid (Fc), resulting in a higher (i.e., “brighter”) or lower (i.e., “duller”) timbre percept, respectively. Increases or decreases in the spacing between harmonics (yellow) correspond to increases or decreases in the F0 of the complex, resulting in a higher or lower pitch percept, respectively. All stimuli were lowpass filtered at 10 kHz. (C) Spectra of all nine complex tones used in the timbre condition (F0 fixed at 200 Hz). (D) Spectra of all nine complex tones used in the pitch condition (Fc fixed at 2400 Hz).

### Topographic Mapping of Both Spectral Content and Fundamental Frequency

To assess the patterns of topographic cortical mapping for each of the three conditions (pure-tone, timbre, and pitch), we constructed a separate feature tuning model, where the general linear model (GLM) beta estimates for each voxel in the region of interest (ROI) in auditory cortices of each participant were characterized as a Gaussian filter applied to the respective stimulus feature (frequency, Fc, or F0). Note that while the analyses were performed on vertices in cortical surface space, the term “voxel” will be used throughout. Figure 2 shows the resulting filters’ center frequencies (CF) in the pure-tone condition on a cortical surface for one representative participant as well as the group average. Both individual and group levels of analysis show robust high-low-high tonotopic gradient reversals, in line with earlier studies (Formisano et al., 2003; Langers and van Dijk, 2012; Thomas et al., 2015), with a region of lower CFs (warmer colors) being anteriorly and posteriorly flanked by regions of higher CFs (cooler colors), centered roughly on HG. At both the individual and group levels, there are additional smaller clusters of low- and high-CF voxels, as reported in earlier studies (e.g., Da Costa et al., 2011b; Moerel et al., 2013).

**Figure 2.**
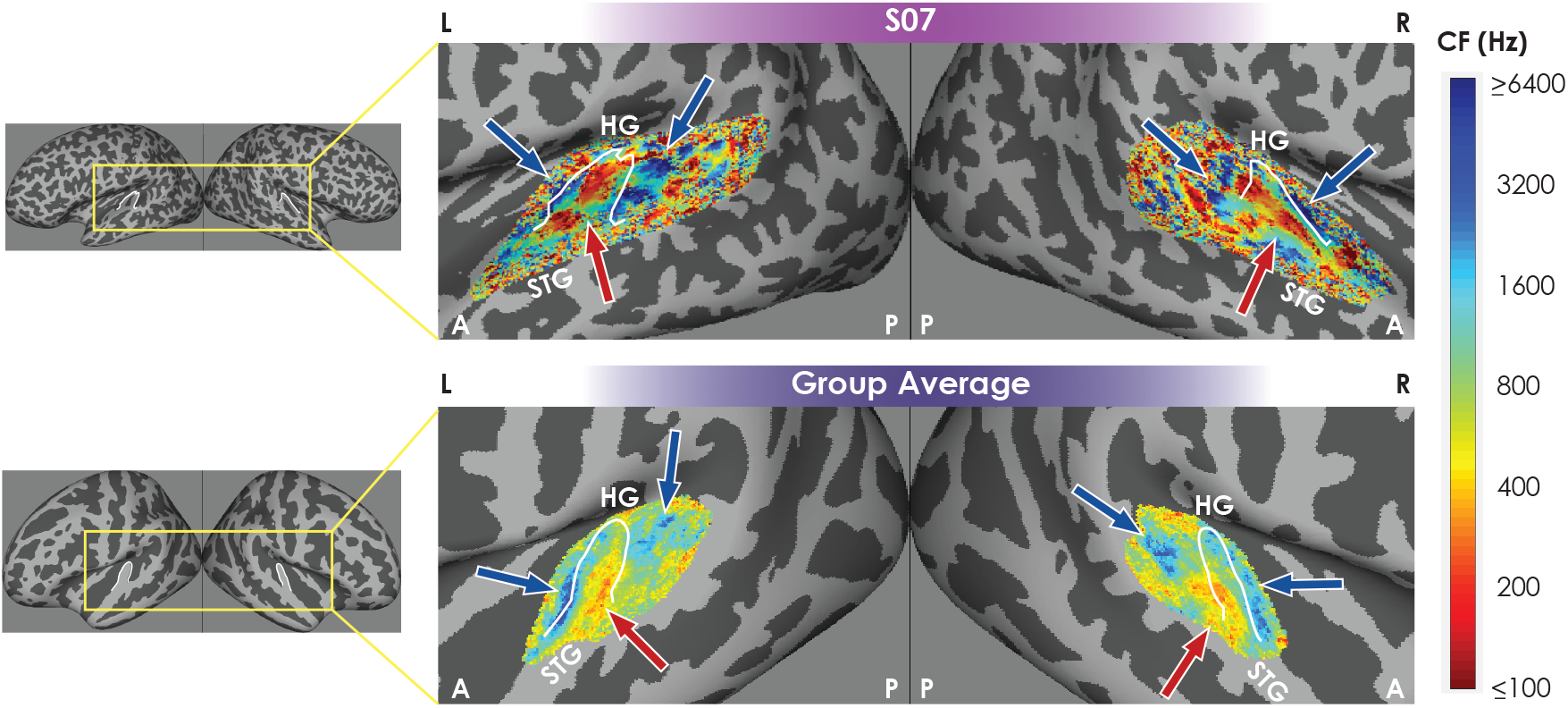
Observations of Tonotopy Elicited by Pure Tones. Feature tuning model CF parameter maps for the pure-tone condition shown on inflated brains for one participant (top) and averaged in fsaverage space across all ten participants (bottom) within the intersection of all participants’ ROIs. Heschl’s gyri denoted by white lines. Blue arrows indicate high frequency regions. Red arrows indicate low frequency regions. L = left hemisphere, R = right hemisphere, A = anterior, P = posterior, HG = Heschl’s gyrus, STG = superior temporal gyrus.

To determine whether the well-established tonotopic organization found with pure tones reflects spectral energy or fundamental frequency in more complex sounds, we compared the pure-tone CF maps to the CF maps in the timbre and pitch conditions. Since it can be difficult to visualize the auditory cortices within the Sylvian Fissure on an inflated lateral surface, Figure 3 shows these maps on inflated spherical representations of the cortices for several representative individual subjects and the group average. We found the timbre maps to be broadly similar to the pure-tone maps in terms of their high-low-high (blue-red-blue) structure. Differences between the maps may be partly due to the different frequency ranges tested, with pure tones CFs spanning 100-6400 Hz and timbre tones CFs spanning 400-6400 Hz. The topographic organization in the pitch condition seems less well defined, although a similar high-low-high gradient can be identified in both the individual and group-average data. The cortical locations of the high and low CF regions are reasonably similar for the timbre and pitch maps, despite the fact that they are derived from independent acoustic features – the spectral peak and the F0, respectively. The individual maps illustrate substantial inter-subject variability in anatomy and function. For instance, S03 has two HG in each hemisphere, and the low CF region is not centered on HG, but instead falls between the two HG. Maps for each of the ten participants are shown in the Supplementary Material, Figure S1.

**Figure 3.**
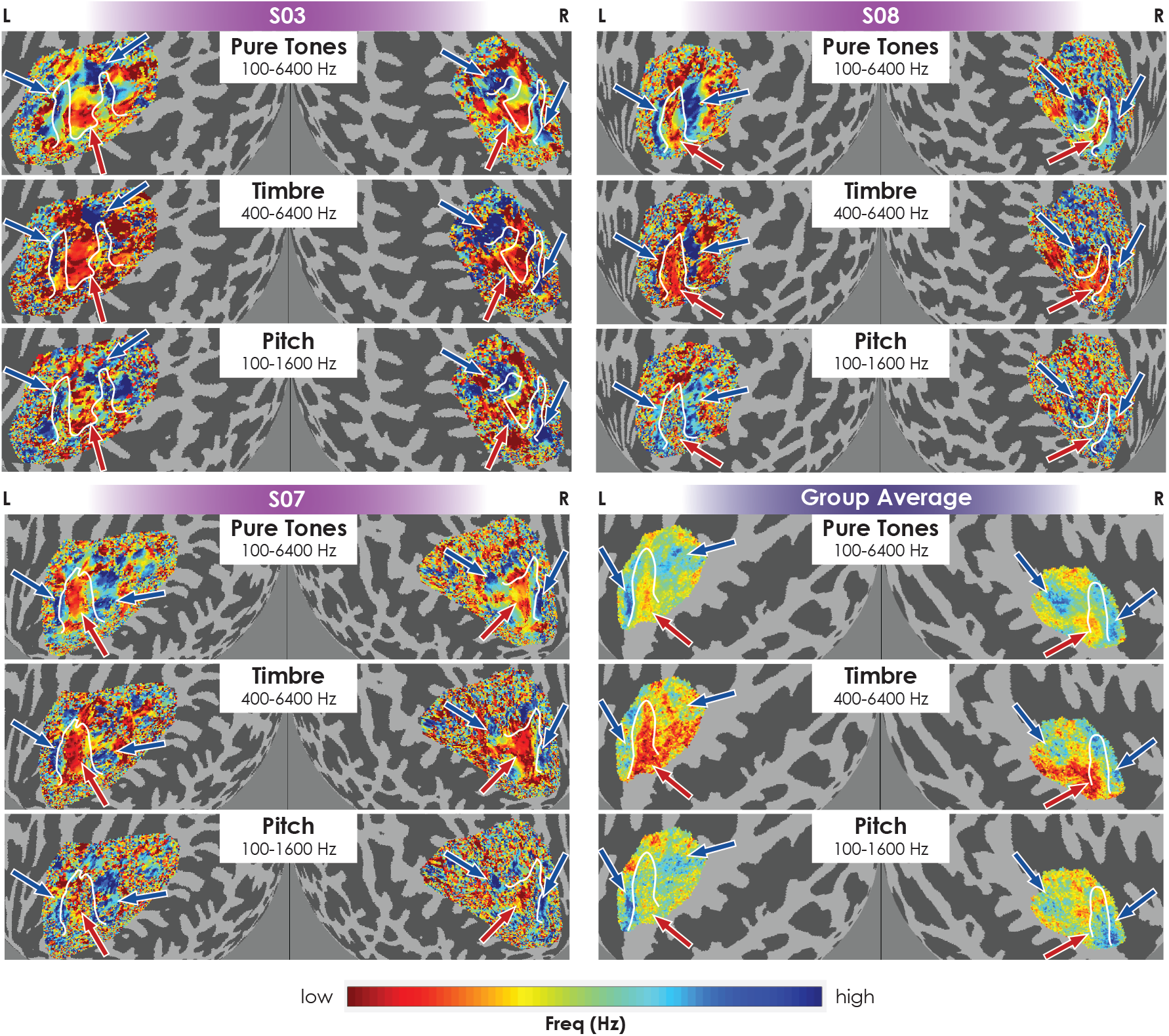
CF Maps for All Three Conditions Show Gradient Reversals. Unthresholded CF maps for the Feature Tuning Model for three individual participants and averaged across all participants (bottom right quadrant). Maps are shown on inflated cortical spheres of the left and right hemispheres, respectively. Each row is the CF map for a given condition (pure tones, timbre, and pitch, respectively). Blue and red arrows indicate high- and low-CF regions, respectively, in the pure-tone conditions. Arrows are shown in the same anatomical locations for the timbre and pitch conditions for ease of reference. Custom color maps span the respective CF range, on a log scale, of each condition, as labeled. L = left hemisphere, R = right hemisphere. HG denoted by white lines.

Bandwidths (BW) of the Gaussian filters were also estimated for each condition. For both the pure-tone and timbre conditions, the narrowest bandwidths tend to be clustered centrally, around HG, consistent with earlier reports using just pure tones (Thomas et al., 2015). The distribution of BWs for the pitch condition is again less clear cut, although some participants show some indication of a central region with sharper tuning. See Figure S2 maps of the BW estimates for all ten participants.

To better understand which regions in auditory cortex are driven by each condition, Figure 4 shows the variance accounted for (R^2^) in the beta weights by each voxel’s filter in each of the three conditions. As with the model CF and BW parameters, the spatial distribution of the high R^2^ voxels is similar in the pure-tone and timbre conditions. In the pitch condition, the number of voxels with a substantial amount of variance explained is reduced, with the exception of regions surrounding HG. While there appears to be some inter-individual variability in the spatial patterns of high R^2^ voxels, the group average map shows a small cluster of higher R^2^ values lining the anterolateral side of HG, bilaterally (Figure 4, lower right). This is consistent with previous studies’ reports of the location of pitch-sensitive regions in both humans (Norman-Haignere et al., 2013; Penagos et al., 2004) and non-human primates (Bendor and Wang, 2006). The R^2^ maps for each of the ten participants are shown in Figure S3, and a comparison of data and feature tuning model fits is provided in Figure S4.

**Figure 4.**
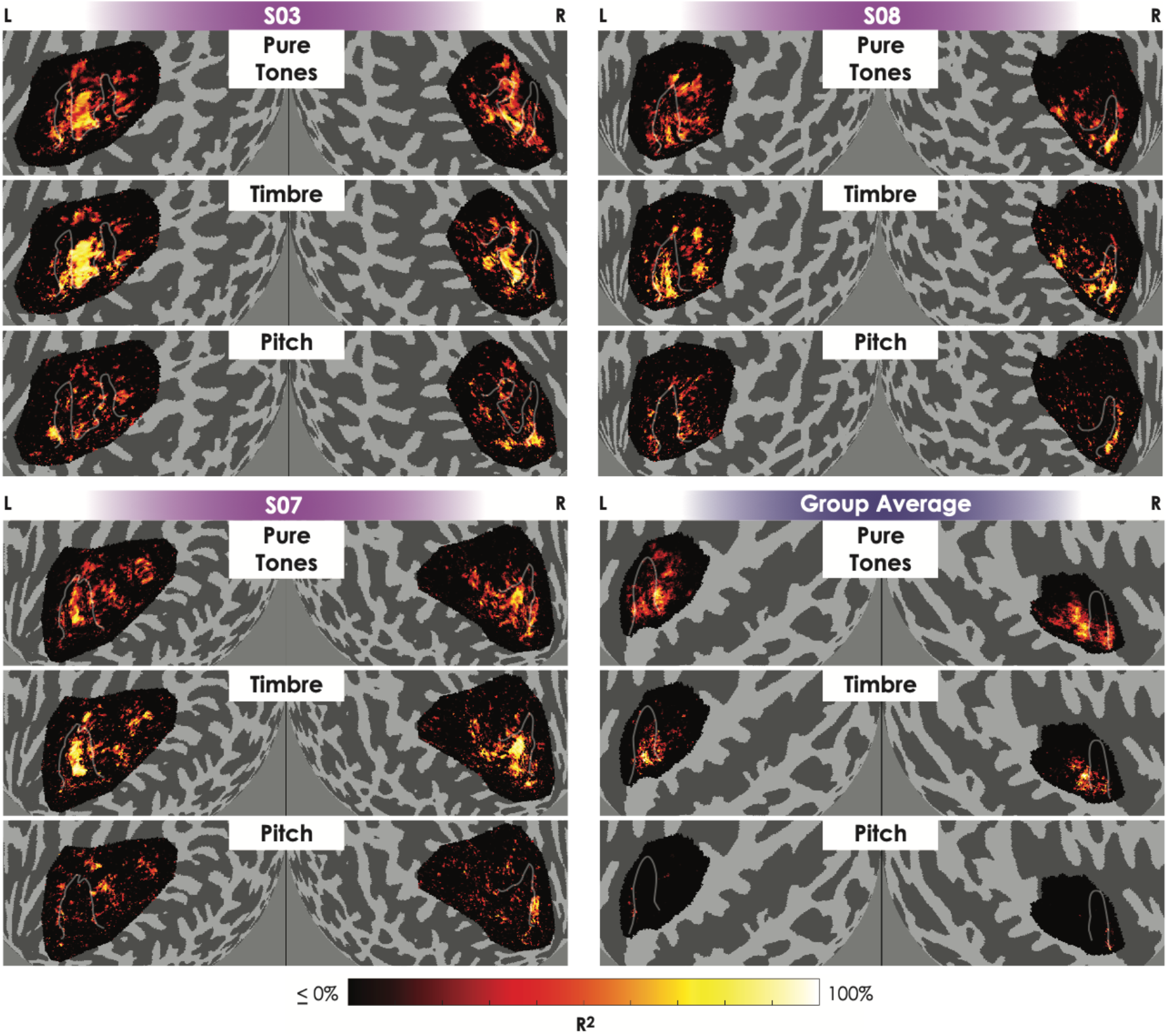
Variance Explained Heat Maps are More Similar for Pure-tone and Timbre Conditions. R^2^ heat maps for the feature tuning model for three participants and averaged across all ten participants (bottom right quadrant) on inflated cortical spheres of the left and right hemispheres, respectively. Each row is a different condition. L = left hemisphere, R = right hemisphere. HG denoted by white lines.

### Pure-tone Cortical Tonotopy Primarily Reflects Spectral Content

The analysis shown in Figures 3 and 4 for each condition separately indicates strong similarities between the pure-tone and timbre conditions, suggesting that traditional pure-tone tonotopy measures primarily reflect a sound’s spectral content, rather than its pitch. To provide a more direct comparison of the cortical responses for different conditions, we calculated representational similarity matrices (RSMs) both within and across conditions (Figure 5). Each matrix cell shows the correlation coefficient between voxels’ responses for a given pair of stimuli (in terms of F, Fc, or F0) within the ROI. High correlations in cells near the main diagonal, as seen in the within-condition comparisons for both pure-tone and timbre conditions (upper-left and center boxes in each panel), indicate that tones that are closer in frequency (or Fc or F0) produce activation patterns that are more strongly correlated across voxels than tones that are distant in frequency. A similar diagonal correlation pattern can be seen when comparing patterns of activation *between* the pure-tone and timbre conditions (upper-center box), suggesting that voxels are responding to similar features in both conditions. In contrast, within-condition comparisons for the pitch condition (bottom right box) show higher correlations across all tones, and the RSMs comparing pitch and pure-tone conditions and comparing pitch and timbre conditions (upper- and center-right boxes) show similarly high correlations for all higher frequencies (or Fcs), independent of F0. This is likely driven by the relatively high (2400 Hz) spectral center of all pitch stimuli. Overall, the RSM analysis confirms our initial analysis showing that classic tonotopy likely reflects the spectral content, and not the F0, of complex sounds. The RSMs for each of the ten participants are shown in Figure S5.

**Figure 5.**
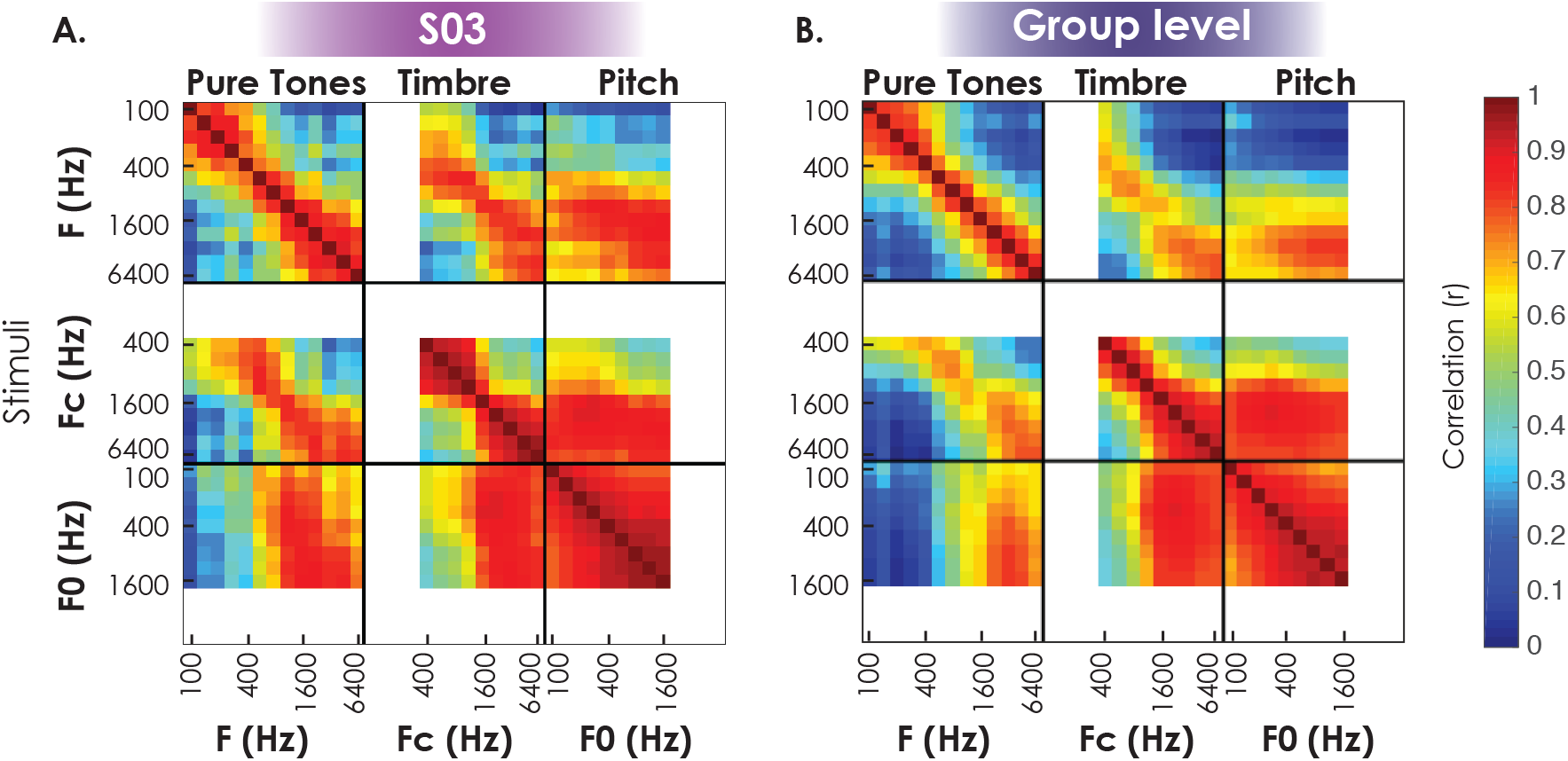
Patterns of Activation are More Similar for Pure-tones and Timbre Conditions. (A) RSMs within the ROI of one participant, averaged across repeats for a given stimulus, thresholded to include voxels with a GLM R^2^ of at least 10% (i.e., voxels that show robust sound-evoked responses). White spaces indicate pure tone frequencies that do not have corresponding Fc or F0 values for the timbre and pitch conditions, respectively. (B) RSMs thresholded at a GLM R^2^ of 10% for each participant and then averaged across all ten participants. Same plotting conventions as in Panel A.

### Similarities in Cortical Tuning Properties for Pure Tones, Timbre and Pitch

Although the pure-tone response patterns seem to resemble those of the timbre condition, reflecting spectral content, some similarities in topographic mapping were also observed between the pure-tone and pitch conditions, reflecting sensitivity to F0. Here we provide a quantitative assessment of these similarities by comparing the model’s CFs obtained in the different conditions for individual voxels. Figure 6 plots the CFs derived from the feature tuning model across pairs of dimensions, including only voxels with overlapping CF ranges for the conditions being compared, and whose predicted responses accounted for at least 30% of the feature tuning model variance across both conditions. As expected, there was a strong relationship between voxel CFs derived in the pure-tone condition and the CFs for the same voxels derived in the timbre condition (r=0.89; N=3256), with an average relationship close to unity.

**Figure 6.**
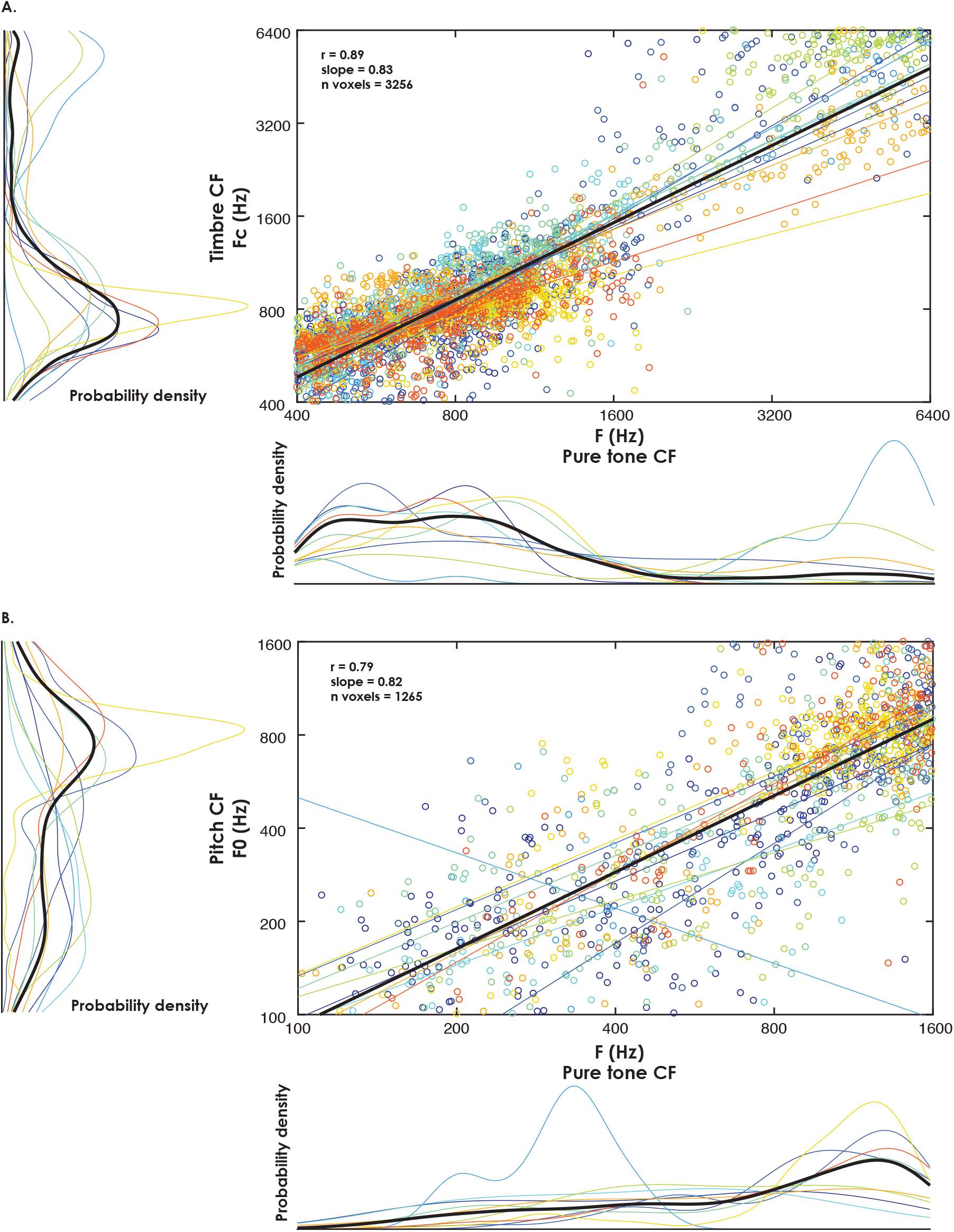
Auditory Cortical Voxels Show Similar Tuning CFs Between Features. (A) Scatterplots comparing CFs for the feature tuning model for timbre and pure-tone conditions. Voxels included for each participant are the intersection of those that exceeded the feature tuning model R^2^ threshold of 30% for each condition being plotted and only for CFs that were included in both conditions being compared. Each participant’s data is a different color. Fit lines for each participant’s data, as well as mean fit lines (black) are plotted. To the left and bottom of each scatterplot are marginal kernel density histograms as well as a mean kernel density line (black). Pearson’s r, slope of the line fit, and number of total voxels (n voxels) are reported. (B) Scatterplots comparing CFs for the feature tuning model for pitch and pure-tone conditions. Same plotting conventions as in Panel A.

Interestingly, although there were fewer voxels that were responsive to both pure-tone and pitch conditions (N=1265), the correlation between the CFs for those voxels was similarly high (r=0.79). This finding suggests that many voxels responsive to pitch are also responsive to pure tones with a best frequency corresponding to the best F0. Finally, the kernel density histograms indicate a broad peak of voxels with CFs (in terms of frequency, Fc, and F0) around 800 Hz, suggesting a somewhat nonuniform distribution of CFs.

### Shared and Distinct Tuning Properties

The previous section concentrated on voxels that demonstrated tuning (i.e., selectivity along the dimension being tested) in at least two conditions. However, other patterns of tuning were also observed, including voxels that showed tuning specific to just one of the conditions. We investigated these properties by categorizing each voxel as being selective along a certain dimension if the fitted Gaussian function for that voxel accounted for at least 30% of the variance in that condition. We did this for each of the three conditions (pure tones, pitch and timbre), resulting in each voxel being categorized independently as selective (or not) along each of the three dimensions. Figure 7A provides a surface map for one participant, with voxels color-coded to indicate the condition(s) under which the voxels were categorized as selective.

**Figure 7.**
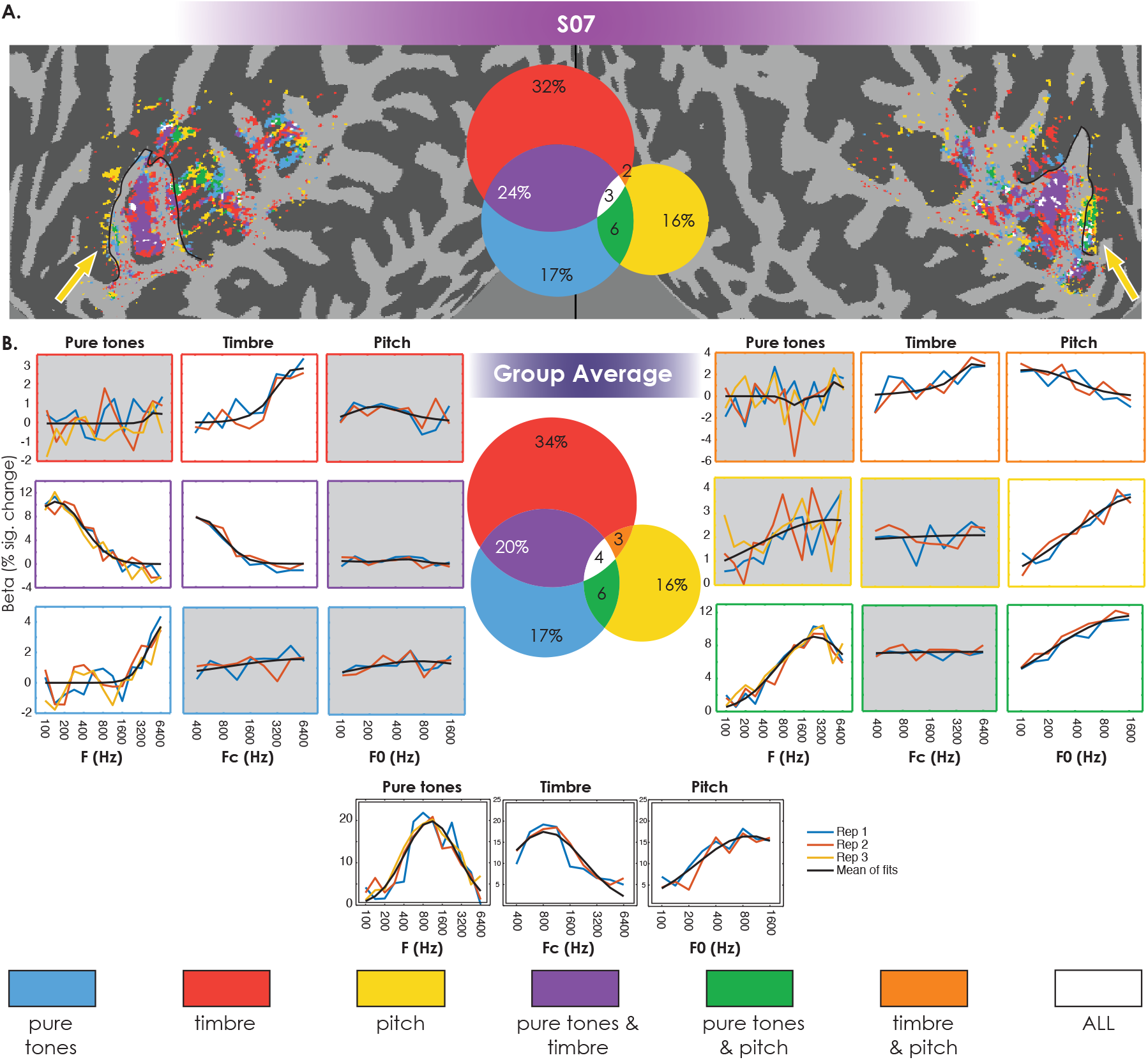
Percentage of Voxels Showing Joint Tuning. (A) Surface map (top) for one participant showing voxels with a model R^2^ of at least 30% for one or more features within the feature tuning model. Different colors denote conditions that meet this threshold, indicating which voxels are demonstrating tuning across multiple conditions. Black outlines denote HG. Yellow arrows point to the regions anterior to HG predominantly tuned to pitch (or pitch and pure tones). In the center is a Venn diagram for this participant showing the percentage of voxels with and without joint tuning, also only including voxels with a model R^2^ of at least 30%. (B) Seven voxels showing various types of tuning. The three plots in each row show tuning of a single voxel to each of the conditions. The colored lines are the beta estimates for each stimulus type, ordered from low to high along the x-axis. The black line is the average fit. Boxes filled with gray indicate poor tuning (feature tuning model R^2^ was 15% or less in all cases) for that condition. The colors of the box outlines correspond to the colors in the Venn diagram. Normalized data from each participant’s individual Venn diagram was averaged together to derive the mean proportions. See legend at the bottom for a description of the color map.

In general, HG contains a large portion of voxels jointly tuned to pure tones and timbre, as well as many voxels tuned specifically to timbre. Beyond HG, while all combinations of tuning are represented in regions posterior to HG, there are prominent clusters of voxels in regions anterior to HG in both hemispheres, either tuned to both pure tones and pitch or just pure tones, in line with the aforementioned pitch sensitive regions. Along with the surface plots in Figure 7A is a Venn diagram showing the proportions of voxels with each type of tuning for the same sample participant. The Venn diagram for the group-average data is shown in Figure 7B, along with examples of the data and model fits from individual voxels that provide examples of selectivity along one, two, or all three dimensions. Surface plots and Venn diagrams for each participant can be found in the Supplementary Material (Figure S6). The relative proportions shown in these Venn diagrams remain similar for a range of R^2^ thresholds and are not specific to the 30% threshold that was chosen (see Figure S7). Overall, the greatest percentage of voxels are tuned to just the timbre of complex tones, followed by voxels jointly tuned to timbre and pure tones. Thus, over 70% of voxels are tuned to some aspect of spectral content (pure-tone, timbre, or both), with a relative lack of tuning to F0. The fact that many voxels appear to have selectivity for the spectral content of complex tones but not for the pure tones is consistent with findings from single-unit studies that have reported many cortical neurons that respond more strongly to spectrally complex sounds than to pure tones (Bendor and Wang, 2005; Feng and Wang, 2017; Rauschecker et al., 1995). Nevertheless, over 20% of voxels appear to have pure-tone frequency tuning without showing similar selectivity for the overall spectral shape of complex sounds.

Although the population appears to be dominated by voxels with spectral content selectivity, the Venn diagram is consistent with our other measures (e.g., Figure 4) in showing a substantial proportion of voxels, approaching 30%, that appear to have F0 tuning, either exclusively or in combination with tuning to other dimensions.

### Pitch Mapping in Auditory Cortex

Although the primary cortical tonotopic gradients seem to be dominated by spectral content, as shown by the close correspondence between responses in the pure-tone and timbre conditions, evidence for tuning to F0 or pitch was also observed in all participants. Voxels from one participant showing tuning to low, medium, and high F0s are shown in Figure 8A. About 16% of voxels demonstrate tuning to F0 but not to pure-tone frequency or spectral centroid. To further explore the spatial mapping of F0, we examined pitch-tuned voxels that were *not* sensitive to changes in either the pure-tone or timbre conditions (i.e., the yellow area in the Venn diagrams). Figure 8B shows a map containing voxels pooled across all subjects with an R^2^ threshold of 0% and a second, more stringent, map with an R^2^ threshold of 30%. These results demonstrate clear topography for exclusively pitch-tuned voxels that is relatively insensitive to the R^2^ cutoff used in the analysis. There is a trend for a high-low-high F0 mapping around the edges of HG, as denoted by the blue and red arrows. This trend appears to be strongly bilateral and, in fact, no significant differences were found for the number of voxels per hemisphere across subjects (p = 0.58). In addition, there appears to be a region along the STG that is tuned predominantly to low F0s, as indicated by the orange arrows. This area has been identified as a pitch-sensitive region (e.g., Norman-Haignere et al., 2013), in addition to the region anterolateral to HG, which contains a large cluster of high F0-tuned voxels.

**Figure 8.**
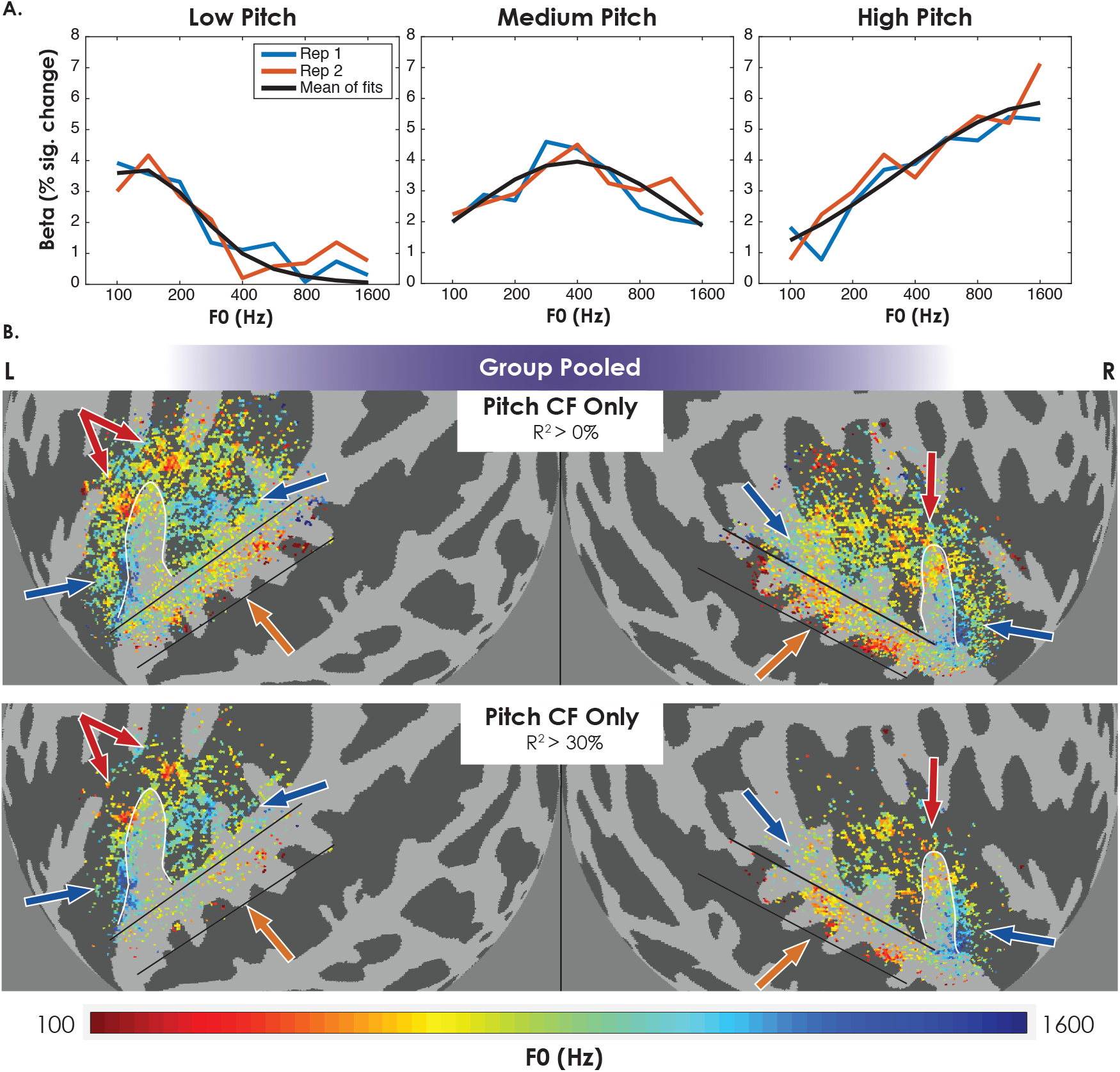
Voxels Showing Clear Pitch Tuning. (A) Three different voxels for one participant tuned to low, medium, and high pitches, respectively. The blue and red lines are the beta estimates, and the black line is the average fit. (B) Voxels pooled across all participants with feature tuning model R^2^ in the pitch condition of > 0% (top) and > 30% (bottom) and excluding voxels with corresponding thresholds in the other two conditions (pure tone, timbre). Voxels in common across participants are averaged. Red arrows point to low-F0 regions and blue arrows point to high-F0 regions. Black lines and orange arrows denote regions along STG tuned predominantly to low F0s. HG denoted by white outlines. For reference, arrows and lines are in the same anatomical locations on both maps.

Critically, the voxels demonstrating clear F0 tuning (R^2^ > 30%) are distributed throughout auditory cortex, bilaterally, and are not located in an isolated region or limited to one hemisphere. This broad distribution may help explain why it has proved difficult to build a consensus on the presence or location of a “pitch region” in auditory cortex (Bendor, 2012; Hall and Plack, 2009). Finally, it is important to note that pitch *sensitivity* (i.e., stronger cortical responses to sounds with greater pitch salience), as has previously been explored, is distinct from pitch *selectivity* (i.e., tuning to certain F0s) that we demonstrate here.

### Spectral Tuning Model

To further support the claim that the topography shown in Figure 8 is a reflection of F0 tuning and cannot be explained by the subtle differences in spectral fine structure that occur with changes in F0, we employed a spectral tuning model. In this model, instead of using the spectral peak as the input for the timbre condition, and F0 as the input for the pitch condition, the Gaussian weighting function was applied to the full sound spectrum and was fitted separately to each of the three conditions. Performance for the pure-tone condition was the same as in the feature tuning model (as the input for both models is the frequency of the pure tone) and performance for the timbre condition was very similar in both models (compare top two rows of Figures 3 and 4 to Figure S8). As expected, given the lack of change in spectral envelope across the range of F0 values tested, this model explained virtually no variance for the pitch conditions (Figure S8B, bottom row), and predicted essentially a flat line across all stimuli in the pitch condition (Figure S8C–E, right panels). The fact that the spectral tuning model could account for the observed responses in the pure-tone and timbre conditions, but not in the pitch condition, further supports the claim that the tonotopy observed in studies using pure tones is predominantly driven by spectral content.

## DISCUSSION

### Summary of Results

This fMRI study attempted to dissociate the auditory cortical mapping of F0 (which determines perceived pitch) from spectral content (which influences timbre) and to determine which of these underlies the well-known tonotopic organization observed with pure tones. Consistent with previous pure-tone studies (Da Costa et al., 2011; Formisano et al., 2003; Saenz and Langers, 2014; Striem-Amit et al., 2011; Thomas et al., 2015), we found bilateral V-shaped high-low-high gradient reversal maps roughly centered on HG in all subjects, as well as narrower tuning bandwidths around HG and broader tuning bandwidths in surrounding regions. Although alignment of subjects is complicated by differences in size, shape, and number of HG in each hemisphere across individuals (Rademacher et al., 2001), this high-low-high pattern of tonotopy was preserved at the group level (Figure 2).

A similar high-low-high pattern of cortical mapping was found with complex sounds that maintained a constant F0 but varied systematically in their spectral peak or centroid, resulting in changes in timbre along a dull-bright continuum (Figure 3). The similarity of maps derived from the pure-tone and timbre conditions both within and beyond HG suggests that the tonotopic maps observed in earlier studies are driven primarily by spectral content and not by F0 or pitch. However, the fact that many voxels exhibited selectivity for spectral content with complex tones but not pure tones and *vice versa* (Figure 7) suggests an organization more complex than simple quasi-linear filtering. Most strikingly, we observed a systematic mapping of pitch, particularly in regions surrounding HG, when examining responses to complex stimuli with a fixed spectral peak but varying in F0 (Figures 3 and 8).

### Relationship to Previous Studies

In addition to pure tones commonly being used to study cortical tonotopy, recordings of complex natural sounds such as speech, musical instruments, and animal vocalizations, have been used to derive feature representations in auditory cortex using fMRI (e.g., De Angelis et al., 2018; Moerel et al., 2012). However, as with pure tones, the positive correlation between F0 and spectral energy often found in natural sounds (Assmann and Nearey, 2008; Hillenbrand and Clark, 2009; McAdams, 2013) makes it difficult to conclude whether the derived maps reflect spectral energy distributions, F0, or a combination of the two. The present study resolves this issue by independently varying F0 and Fc in order to tease apart the cortical topography of these features.

While not explored in fMRI, early MEG studies using complex tones did attempt to study the relationship between pitch and sound spectra. However, they came to differing conclusions, suggesting either that cortical tonotopy reflected pitch, rather than spectral distribution (Pantev et al., 1989), or that it reflected orthogonal representations of both pitch and spectral distribution (Langner et al., 1997). However, the limited spatial resolution of MEG makes it poorly suited to fine-grained analysis of the topographical organization of cortical representations. The present study utilized high field fMRI to explore the topography of these features at a much higher spatial resolution. Although many studies have explored fMRI correlates of general pitch responsivity in human auditory cortex (e.g., De Angelis et al., 2018b; Hall and Plack, 2009; Norman-Haignere et al., 2013; Penagos et al., 2004), the present findings provide new insights into how representations of F0 are organized in the brain and their relationship with spectral content.

### Voxel Tuning to Multiple Dimensions

This study reveals that a considerable proportion of voxels exhibit tuning for two or more of the conditions tested (~33%), particularly for the pure-tone and timbre conditions (Figure 7). However, more voxels (~67%) demonstrated clear tuning properties in only one of the conditions. Although a majority of the voxels that exhibited tuning to more than one dimension were selective to the pure-tone and timbre conditions, it may seem surprising that the overall proportion (~20%) was not greater, given the evidence that tuning in both those conditions are driven by spectral content. This apparent discrepancy may be due in part to the fact that the range of pure-tone frequencies (100-6400 Hz) was greater than the Fc range (400-6400 Hz), but may also reflect genuine differences in selectivity based on higher-level features, rather than just spectral shape. Indeed, our findings are in line with single- and multi-unit studies in other species that have identified neurons that are uniquely sensitive to either pure tones or complex tones, but not both (e.g., Feng and Wang, 2017; Rauschecker et al., 1995). In fact, in marmosets, Bendor and Wang (2005) found a similar percentage of neurons that were tuned to narrowband and broadband complex stimuli that did not respond significantly to pure tones (~38%) as we found voxels tuned to complex timbre tones and not pure tones (~34%).

Our data support the existence of two distinct maps: one organized by frequency selectivity (i.e., tonotopy), and the other organized by pitch selectivity. While these maps are partially overlapping, responses in HG, which is the macroanatomical landmark most closely linked to primary auditory cortex (A1) were predominantly driven by spectral content, as reflected by the strong model fits in both the pure-tone and timbre conditions. Pitch representations, on the other hand, were mostly found in the surrounding non-primary regions. This cortical arrangement is consistent with hierarchical processing of sound, with lower-level frequency content being processed predominantly in A1 and higher-level sound features (e.g., pitch) being processed predominantly in surrounding non-primary (belt and parabelt) regions.

Finally, we observed a non-uniform distribution of voxel CFs (Figure 6), with a larger proportion of voxels being tuned to the frequency region around 800 Hz, which is known to contain the most energy for human speech (Byrne et al., 1994; Cox and Moore, 1988). This organization hints at a possible enhanced representation of acoustic frequencies that are most common and important in our daily lives.

### Pitch Tuning in Auditory Cortex

While many studies have explored responses to pitch in auditory cortex, the approach has generally been to compare sounds with salient pitches to those with weak pitches to identify pitch-sensitive regions (e.g., Hall and Plack, 2009; Norman-Haignere et al., 2013; Penagos et al., 2004). In contrast, the present study explored voxel-wise tuning to different pitches while controlling for spectral variations. We were able to identify voxels in all subjects that exhibited selectivity along the F0 dimension. While our initial CF maps at both the individual and group level suggested a (somewhat noisy) high-low-high mapping of F0, we expanded on this and mapped out voxels in auditory cortex exclusively tuned to stimuli in the pitch condition, while excluding voxels that also exhibited strong tuning (R^2^ > 30%) in the pure-tone and/or timbre conditions (Figure 8). These maps revealed regions distinctly tuned to low, medium, and high F0 CFs distributed throughout auditory cortex. Specifically, bilaterally, there were clear clusters of voxels tuned to low F0s around the medial portion of HG and lining STG, and clear clusters of voxels tuned to high F0s in regions anterolateral to HG.

### Bilaterality in Cortical Representations

For the pure tones, timbre, and pitch conditions, topographic mapping was relatively symmetric across hemispheres for all participants. While this bilaterality in cortical representations has been shown in other studies (e.g., Allen et al., 2017, 2018; De Angelis et al., 2017; Hall and Plack, 2009; Norman-Haignere et al., 2013; Patterson et al., 2002; Penagos et al., 2004; Warren et al., 2005), there is some evidence to suggest a right hemisphere lateralization for some forms of pitch processing (e.g., Albouy et al., 2020; Hyde et al., 2008; Zatorre et al., 2002). However, the present study found no significant difference in the number of pitch-tuned voxels between hemispheres, which suggests that pitch *selectivity* is represented bilaterally.

### Open Questions

With the advancement of methods for measuring fMRI responses from distinct cortical layers (Ahveninen et al., 2016; Kay et al., 2019; Moerel et al., 2020), an interesting topic for future research is whether topographic maps of frequency and pitch have differential signatures as a function of cortical depth. For example, frequency mapping inherited from the cochlea may originate in middle layers that receive thalamocortical connections, whereas more complex processing of perceptual features like pitch may emerge in more superficial layers. Consistent with hierarchical processing of sound across cortical depths, Moerel et al., (2019), used 7T fMRI to demonstrate that a simple frequency model could more accurately characterize responses in deep and middle layers well, while responses in superficial layers were better predicted by features from a more complex spectrotemporal modulation model.

Finally, while the present study found systematic maps of absolute pitch, it remains unclear how relative changes are represented in cortex. Absolute pitch refers to the exact pitch (F0) of a sound (e.g., the musical note A4 is 440 Hz), whereas relative pitch relates to contour and interval size (i.e., whether the pitch is going up or down compared to other pitches and the magnitude of this change). Relative pitch processing is essential for both music and speech comprehension and reflects a higher-order process than absolute F0 encoding. Recent studies, based on recordings from subdurally implanted electrodes, as participants listened to variable pitch contours in speech stimuli, provide evidence of both absolute and relative pitch encoding in human auditory cortex (Hamilton et al., 2020; Tang et al., 2017). Extending upon this work using high-resolution fMRI could provide a more comprehensive characterization of relative tuning throughout auditory cortex and elucidate its relationship to other processes within the auditory hierarchy.

## METHODS

Detailed methods are provided and include the following:
 
- LEAD CONTACT AND MATERIALS AVAILABILITY
- EXPERIMENTAL MODEL AND PARTICIPANT DETAILS
- METHOD DETAILS
- QUANTIFICATION AND STATISTICAL ANALYSIS

## ACKNOWLEDGEMENTS

This work was supported by the National Institute on Deafness and other Communication Disorders (NIDCD, grant number R01 DC005216). The Center for Magnetic Resonance Research (CMRR) is supported by NIH grants P41 EB027061, NIH P30 NS076408, NIH S10 RR026783, and by the W.M. Keck Foundation. Anahita Mehta, Andrea Grant, and Cheryl Olman provided helpful advice, assistance, and training.

## AUTHOR CONTRIBUTIONS

All authors designed the experiment. A.J.O. acquired funding. A.J.O. and K.N.K. provided supervision and resources. E.J.A. and J.M. collected the data. E.J.A. and J.M. analyzed the data. E.J.A. wrote the manuscript. All authors edited the manuscript.

## METHODS

### LEAD CONTACT AND MATERIALS AVAILABILITY

Further information and requests for resources should be directed to and will be fulfilled by the Lead Contact, Emily Allen.

### EXPERIMENTAL MODEL AND PARTICIPANT DETAILS

#### Participants

The Institutional Review Board (IRB) for human participant research at the University of Minnesota approved the experimental procedures. Written and informed consent was obtained from each participant prior to data collection. Ten people from the University of Minnesota community (average [SD] age of 29.3 [4.2] years; 6 females, 4 males), all right-handed and having normal hearing, defined as audiometric pure tone thresholds of 20 dB hearing level (HL) or better, at octave frequencies between 250 Hz and 8 kHz, participated in this study. An eleventh participant was excluded after having great difficulty hearing the stimuli and discovering elevated thresholds since their last audiogram, making them no longer eligible for participation.

### METHOD DETAILS

#### Stimuli and Procedure

All stimuli were generated in MATLAB (The MathWorks) and presented using the Psychophysics Toolbox (Kleiner et al., 2007). Stimuli consisted of three condition types: pure tones, complex pitch tones, and complex timbre tones. The 13 pure tones, each a single frequency, spanned six octaves (100-6400 Hz), in half octave steps (Figure 1A). The pitch and timbre tones were band-pass filtered harmonic complexes (Figure 1B). All complex tones had a 12 dB per octave slope around the center frequency, and a 16^th^ order lowpass filter cutoff at 10 kHz. The nine complex timbre tones had a fixed F0 of 200 Hz and varied in the location of their spectral envelope peak in the frequency domain, or spectral centroid, spanning four octaves (400-6400 Hz) in half octave steps (Figure 1C). The nine complex pitch tones had a fixed spectral envelope centered on 2400 Hz and a varying F0, which spanned four octaves (100-1600 Hz) in half octave steps (Figure 1D). The ranges for the pitch and timbre conditions were chosen to ensure that the spectral centroid was well above the F0 in order to be defined by the stimuli.

Stimuli were presented via MRI-compatible Sensimetrics S14 foam tip earbuds with custom filters to flatten the frequency response (Malden, MA). Stimuli were adjusted to be of equal perceptual loudness. This was done by having two participants, in a separate session, listen to repeats of a single tone type, in blocks lasting 15 s, as the level steadily increased. Participants were then instructed to adjust the level until the tone was clearly and comfortably audible by pressing button “1” on the button box to decrease the level and button “2” to increase the level. Pressing “3” meant they were satisfied with the current level and could advance to the next block. If they did not press “3”, they would automatically advance to the next block at the end of the 15-s block. In each subsequent block they were instructed to make the tone similarly audible as the level chosen in the previous block. The participants performed three repetitions of this task while wearing the Sensimetrics S14 earphones, with the aim of making all sounds equal in level. We then took the median level of these trials and increased all tones by 25-30 dB to be easily heard over the scanner noise. Since equal loudness percepts across frequencies tend to compress at higher levels (ISO, 2003), and since people are differentially affected by background noise, these levels were further adjusted and customized for each participant prior to being scanned, and tweaked at the beginning of the session, in the presence of the scanner noise, as needed. The equal loudness contours were similar across participants, with only small offsets in the mean level required for comfortable audibility. The mean [SD] level (dB SPL) was 83.4 [5.2] for the pure tones, 75.3 [1.8] for the pitch tones, and 80.2 [3.9] for the timbre tones.

We incorporated a “Morse code”-like rhythm into the stimuli designed for this study in order to enhance their perceptual salience over the sound of the magnetic resonance (MR) pulse sequence, inspired by the stimulus design of Thomas et al. (2015a). Each stimulus was presented with an equal number of short (50 ms) and long (200 ms) tone bursts, including 20-ms onset and offset ramps. Every 700 ms consisted of two short and two long tones, pseudorandomly positioned within the 700 ms period, with a 50-ms gap after each tone. This was repeated 11 times for a total stimulus length of 7.7 s. All tones presented within the 7.7 s had an identical F0 and spectral centroid but varied in duration. After a 700-ms gap, a new stimulus was presented (with a new frequency, F0, or spectral centroid, depending on the condition) for 7.7 s, and so on, until all tones of a given condition were presented once in a pseudorandomized order (i.e., one condition block), followed by a 12-s silent gap. There were a total of 12 experimental runs (three pure-tone runs and nine complex-tone runs), each about six minutes long. The order of the pure- and complex-tone runs was counterbalanced across participants. Each pure-tone run consisted of three pure-tone blocks and each complex-tone run consisted of two pitch blocks and two timbre blocks, presented in a pseudorandomized order. Ten seconds of padding was added to the beginning and end of each run. A schematic of runs within a session is shown in Figure S9. Participants were instructed to keep very still and resist any desire to move to the rhythm of the stimuli. Their task was to indicate, via button box, whether the 7.7-s tone sequence was lower or higher (in either pitch or timbral brightness) than the previous sequence.

#### Magnetic Resonance Imaging

All data were acquired using Siemens scanners at the Center for Magnetic Resonance Research (CMRR, University of Minnesota). Functional data were acquired at the passively shielded 7T Siemens MAGNETOM scanner using a single transmit 32-channel Nova Medical head coil. The acquisition parameters for the gradient-echo EPI sequence used were: repetition time (TR) = 1400ms; echo time (TE) = 20ms, multiband factor = 2; generalized autocalibrating partially parallel acquisition (GRAPPA) acceleration factor = 3; number of slices = 44; 1.1mm isotropic voxels. Slices were angled to align with the Sylvian Fissure of each participant in order to fully encapsulate auditory cortices. The sound level of the functional sequence at the center of the bore was 101 dBA SPL. Four fieldmaps were also collected throughout each session for distortion correction. The acquisition parameters for the fieldmaps were: TR = 190ms; first echo time = 4.08 ms; second echo time = 5.1 ms; 2.2mm isotropic voxels; 22 slices. The complex tone runs had 258 volumes and the pure tone runs had 267 volumes.

Anatomical (T1 and T2-weighted) data were acquired at the Siemens 3T Prisma scanner with a 32-channel head coil. MPRAGE T1-weighted parameters were: TR = 2400ms; inversion time (TI) = 1000ms; TE = 2.22ms; flip angle = 8°; 0.8mm isotropic voxels. T2-weighted parameters were: TR = 3200ms; TE = 563ms; 0.8mm isotropic voxels. Six T1s and three T2s were acquired for each participant.

Half of the participants used custom foam Caseforge head cases (caseforge.co). The posterior portion of each head case was used to help stabilize participants’ heads during the scans and additional padding was added under the neck and around the ears for further stabilization and comfort. The other participants used standard MR-compatible foam padding on the back of the head, along with additional neck and ear padding.

### QUANTIFICATION AND STATISTICAL ANALYSIS

#### Anatomical and Functional Preprocessing

The data were preprocessed using a custom pipeline developed by Kendrick Kay’s Computational Visual Neuroscience (CVN) lab (Kay et al., 2019). Gradient unwarping, which corrects image distortions due to gradient nonlinearities, was performed on the T1 and T2-weighted anatomical volumes using the gradient coefficient file provided by Siemens. All six T1 volumes for a given participant were then co-registered using rigid-body transformation with six degrees of freedom and cubic interpolation. Once aligned, the volumes were averaged together to improve contrast between the gray and white matter for high quality segmentation. The same process was used for the three T2 volumes. The averaged T2 volume was then aligned to the averaged T1 volume for each participant.

Cortical reconstruction was performed via FreeSurfer (Fischl, 2012) using the averaged T1 volume. Since the anatomical data had sub-millimeter resolution, a “hires” flag was added, and an expert file was used to specify a larger number of inflation iterations (50). Segmentation results were then visually inspected in Freeview. The functional data was sampled across the cortical thickness at 25%, 50%, and 75% cortical depths and then averaged together. To plot group-level maps, individual subject results were mapped to FreeSurfer’s fsaverage cortical surface group space via nearest-neighbor interpolation.

Functional data preprocessing included slice time correction, fieldmap-based undistortion, and motion correction. Functional data were aligned to the anatomical data using an affine transformation. In the slice time correction step, the data were temporally upsampled from 1.4s to 1s. In the motion correction step, the data were sampled onto the FreeSurfer depth-dependent surfaces. No smoothing was applied to the data.

The data were denoised and GLM analyses were run using the GLMdenoise toolbox (Kay et al., 2013). Each 7.7-s tone sequence was analyzed as a block, and a canonical hemodynamic response function (HRF) was assumed. Leave-one-run-out cross-validation was performed and R^2^ was used to quantify the proportion of the time-series variance (R^2^) that can be explained by the stimuli across all conditions. The three pure tone runs (each containing three repetitions of each tone) were used to estimate three betas per tone. Likewise, the nine complex tone runs (each containing two repetitions of each tone) were used to estimate two betas for each pitch and timbre tone.

#### Encoding Models

Encoding models were used to explore how similar or dissimilar topographic representations of pure tones were to the representations of tones varying either in their F0 or spectral centroid. The first model we used was the feature tuning model. For this model, the following equation was used:

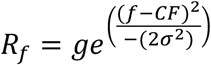

where responses (*R*) to a given feature (*f*) were modelled using a Gaussian function described by three parameters: gain (*g*), center frequency (*CF*), and standard deviation (σ) of the Gaussian. While the architecture of this model was identical across conditions, the input into this model varied. Specifically, for pure tones, a single frequency value was the input for each of the pure tone stimuli, a single F0 value was the input for each of the pitch stimuli, and a single Fc value was the input for each of the timbre stimuli. We assessed model performance using n-fold cross-validation, with pure tones having three folds (two betas per stimulus used for training, one for testing), and pitch and timbre each having two folds (one beta per stimulus for training, one for testing), due to the number of beta estimates that came out of the GLM analysis. For each fold, model R^2^ was derived using the held-out data, by computing the proportion of the original variance in the data that was unaccounted for by the model fit and subtracting this quantity from 1:

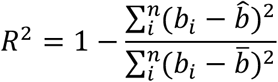

where *b* is the pattern of beta weights across *n* stimuli in a given condition, 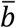 is the mean across beta estimates, and 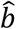 are the predicted betas of the model for the corresponding stimuli. Higher *R*^2^ values indicate more accurate model predictions of the variability in beta estimates across tones of a given conditions.

The second model we implemented was the spectral tuning model, which was inspired by the Population Receptive Field (pRF) method (Dumoulin and Wandell, 2008; Thomas et al., 2015). Instead of characterizing responses to each stimulus on the basis of a single-valued stimulus property, as was done for the feature tuning model, the spectral tuning model took into account the entire frequency spectrum of each stimulus. The form of this model is the same as equation 1, except the response R_-_ depends on the full amplitude spectrum, *S*, of a given stimulus, sampled at frequencies *f* up to 10 kHz:

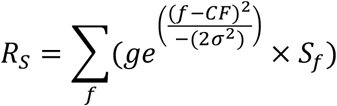

While this model was the same for the pure tone stimuli, which were characterized as a single frequency in both cases, it did change the input for the pitch and timbre stimuli, which are harmonic complex tones containing many frequencies. Because the input feature for this model was frequency spectrum of the stimuli, the same model could be simultaneously applied to all conditions. However, in order to more closely compare the results of the feature tuning model to the spectral tuning model, this model was applied to each condition separately (Figure 9).

#### Regions of Interest

The ROIs for each participant (one per hemisphere) were defined based on several criteria: macroanatomical landmarks of auditory cortices (identifying the Heschl’s gyri for each participant), myelin density maps, and functional data (i.e., the pure tone tonotopy results of the feature tuning model). The macroanatomy served as a starting point for the general location of each region of interest. This was then fine-tuned based on myelin density observed in and around that region. The myelin density maps were generated by dividing the averaged T1 by the aligned and averaged T2 of a given participant. Myelin density was sampled across the cortical thickness at 25%, 50%, and 75% cortical depths. These were then averaged together for a mapping of density across cortical depths. For all participants, these maps showed greatest cortical myelin density in somatosensory, visual, and auditory regions, consistent with earlier studies (e.g., Glasser and Van Essen, 2011). The maps were further refined with the functional data to ensure that the defined ROIs were not too conservative, so as to be missing parts of the tonotopic maps, but also not too liberal, so as to include an excessive number of uninformative voxels. These ROIs were then used across all maps of the modeling results for a given participant.

