## Supplementary Material for "Dissociation of tonotopy and pitch in human auditory cortex"

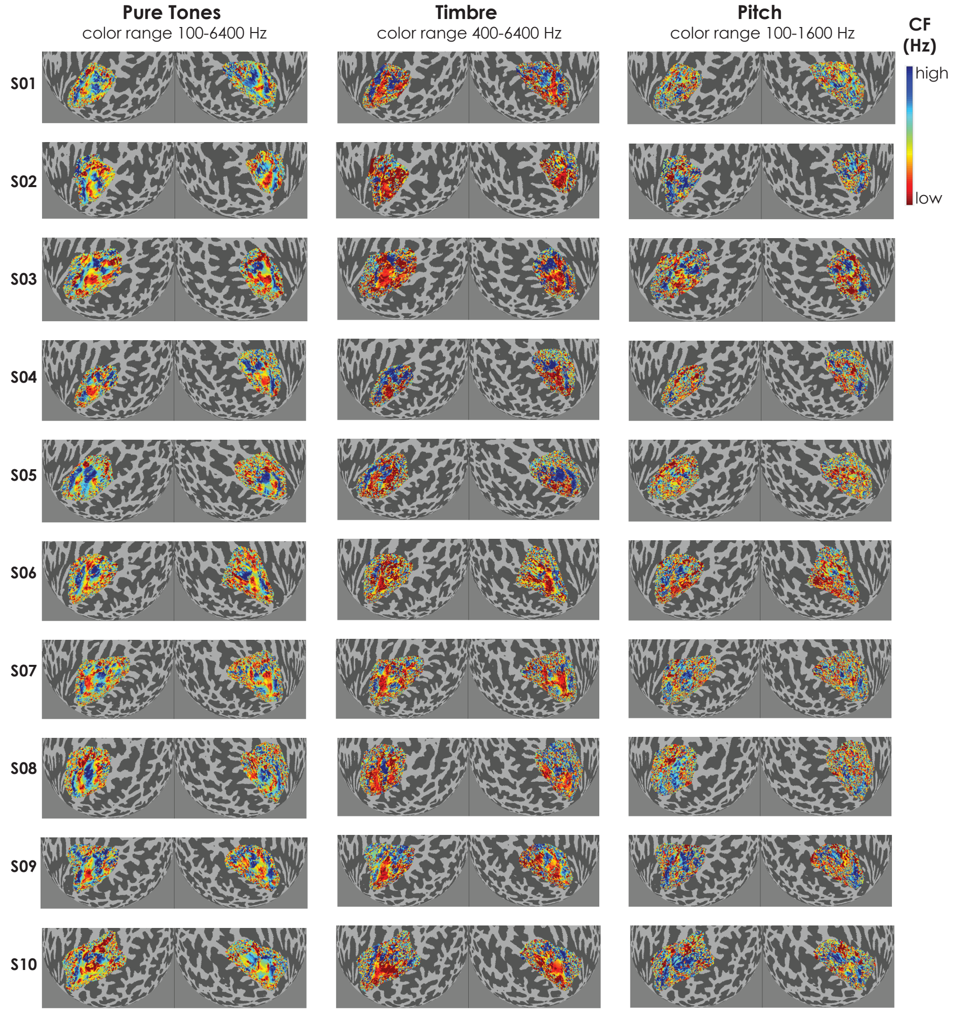


**Figure S1. CF Maps for Each Participant**

Unthresholded center frequency (CF) maps for all ten participants for the feature tuning model within their ROIs. Maps are shown on inflated cortical spheres of the left and right hemispheres, respectively. Each row is a participant. Each column is the CF maps for a given condition (pure tones, timbre, and pitch, representing frequency, center frequency, and fundamental frequency, respectively). Custom color maps span the respective CF range of each condition, as labeled.

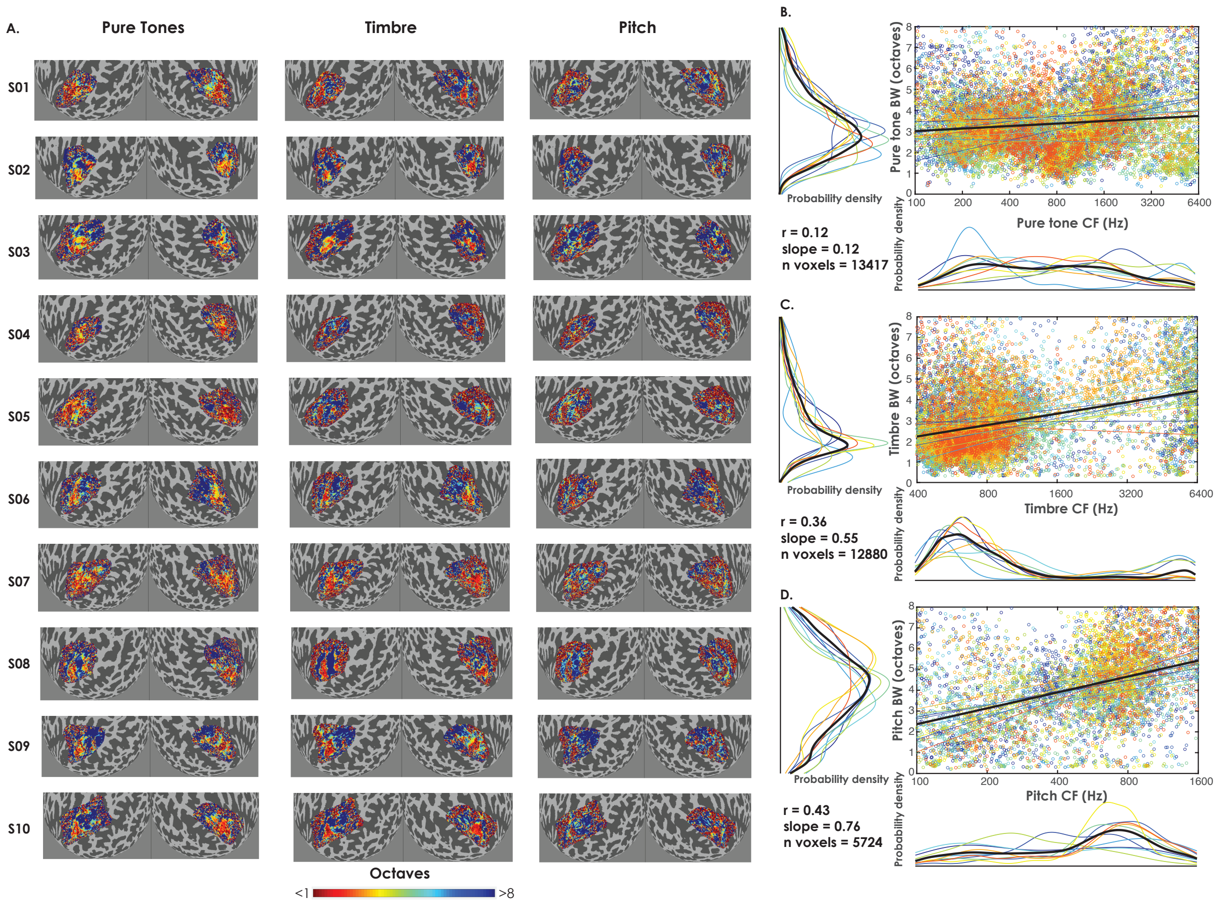


**Figure S2. Distribution of Tuning Center Frequencies and Relationship to Tuning Bandwidths**

(A) Unthresholded bandwidth (BW) maps, defined as the full width at half maximum (FWHM) of the fitted Gaussian (equivalent to 2.355SD), for all ten participants for the feature tuning model within their ROIs. Maps are shown on inflated cortical spheres of the left and right hemispheres, respectively. Each row is a participant. Each column is the BW maps for a given condition (pure tones, timbre, and pitch, respectively). A low number of octaves (reds) indicate a small BW, whereas a high number of octaves (blues) indicate a large BW, corresponding to sharper and broader tuning, respectively.

(B) Scatterplots comparing BWs and CFs for the feature tuning model for the pure tone condition. Voxels included for each participant are those that exceeded the feature tuning model $R^2$ threshold of 30% and had BWs of 8 octaves or less. Each participant's data is shown in a different color. Fit lines for each participant's data, as well as mean fit lines (black) are plotted. To the left and bottom of the scatterplot are marginal kernel density histograms. Pearson’s r, slope of the line fit, and number of total voxels (n voxels) are reported.

(C) Scatterplots comparing BWs and CFs for the timbre condition. Same plotting conventions are used as in Panel B.

(D) Scatterplots comparing BWs and CFs for the pitch condition. Same plotting conventions are used as in Panels B and C.

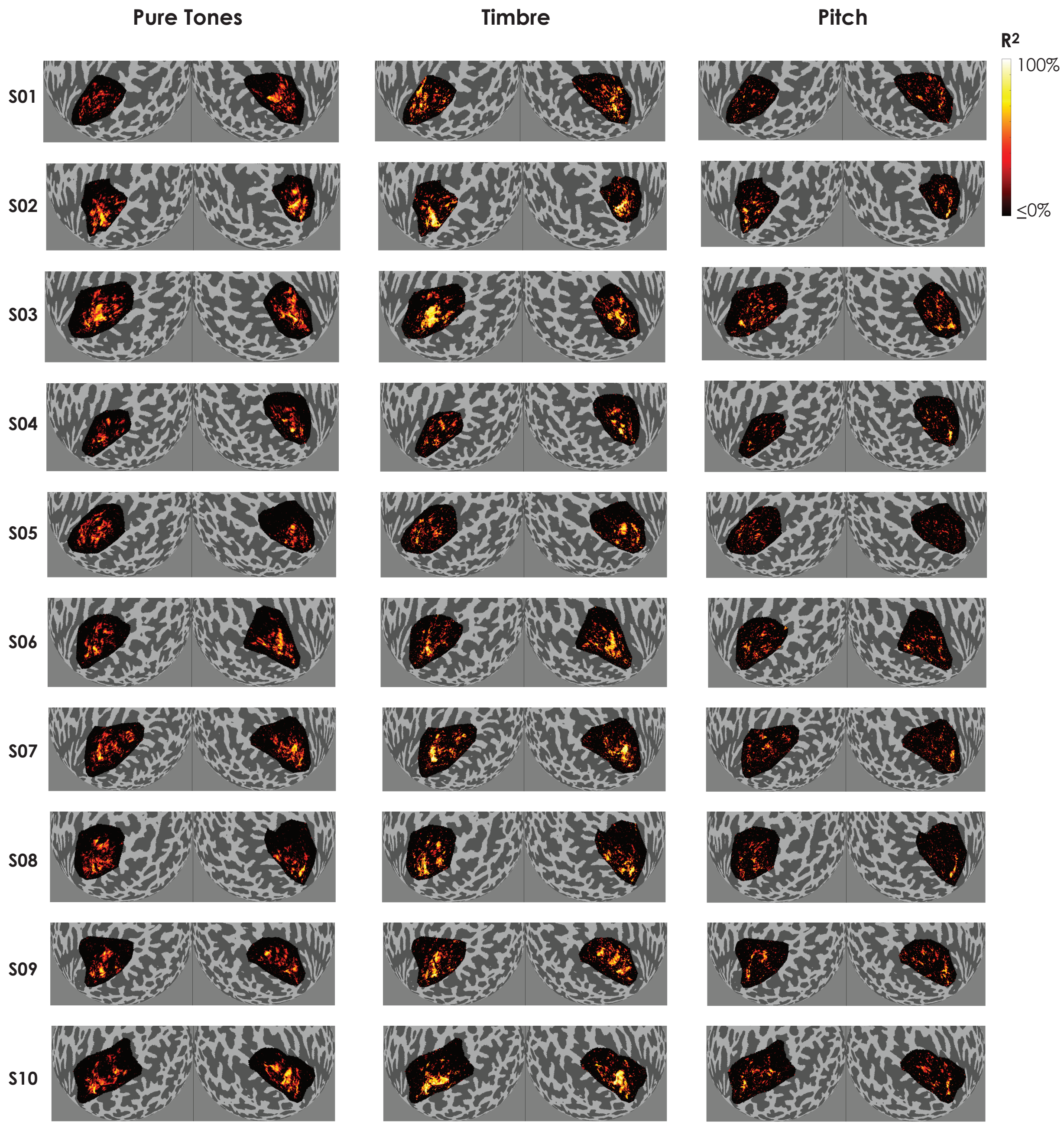


**Figure S3. Maps of Variance Explained for Each Participant**

Heat maps showing $R^2$ values for all ten participants for the feature tuning model on inflated cortical spheres of the left and right hemispheres, respectively. Each row is a participant. Each column is the $R^2$ maps for a given condition (pure tones, timbre, and pitch, respectively).

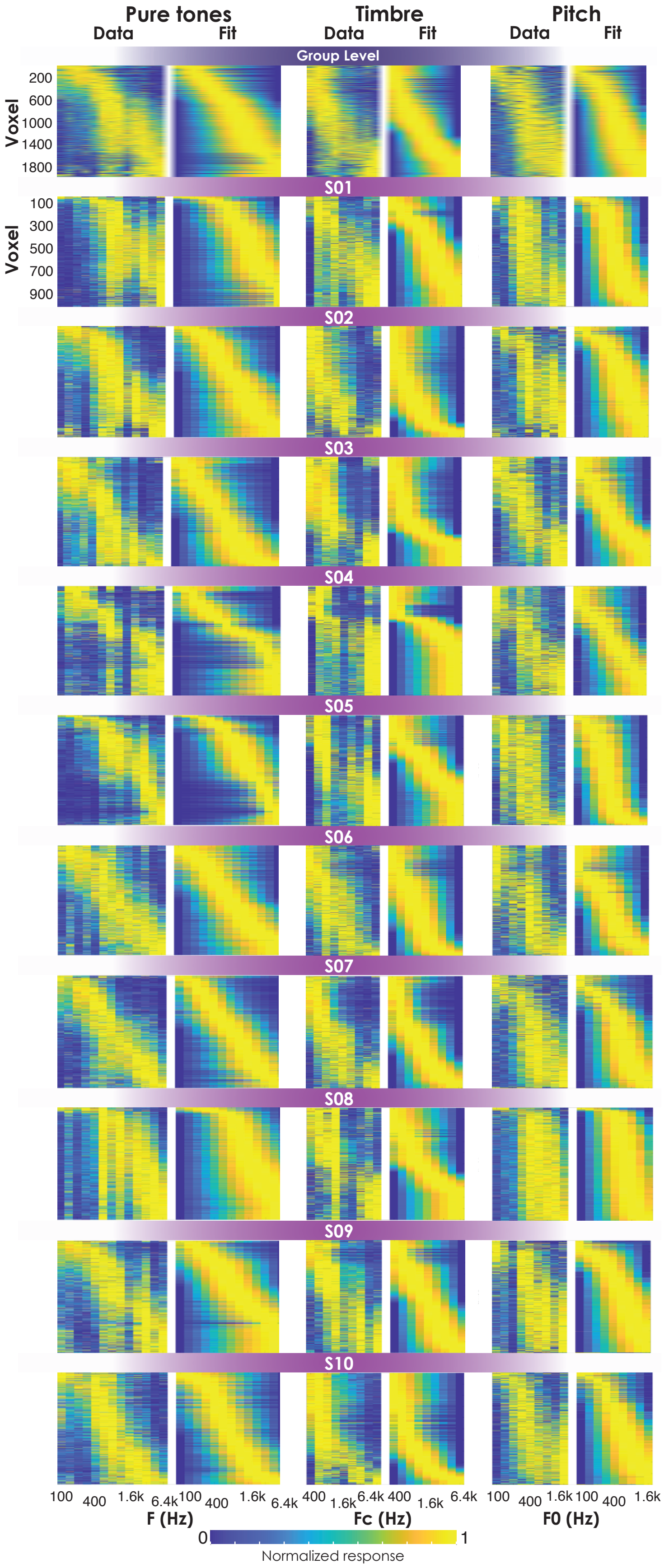

Pure tones
Timbre
Pitch
Data
Fit
Data
Fit
Data
Fit
Group Level
Voxel
200
600
1000
1400
1800
S01
Voxel
100
300
500
700
900
S02
S03
S04
S05
S06
S07
S08
S09
S10
100
400
1.6k
6.4k
100
400
1.6k
6.4k
400
1.6k
6.4k
400
1.6k
6.4k
100
400
1.6k
100
400
1.6k
F (Hz)
Fc (Hz)
F0 (Hz)
0
1
Normalized response

**Figure S4. Comparison of Data and Feature Tuning Model Fits**

Each pair of panels shows the tuning of individual voxels (left) and the predicted tuning of the corresponding model voxel (right) for the voxels with the highest $R^2$ values, ordered from top to bottom based on the fitted CF (from low to high). The three columns show data from the pure-tone, timbre, and pitch conditions, respectively. The top row shows the data pooled across participants, taking only the top 2000 voxels (i.e., voxels with the highest $R^2$ values); the remaining rows show data from each individual participant, taking the top 1000 voxels from each. For visualization purposes, each fit was normalized (0-1 range) and the same scaling constants were applied to that voxel's data. The data show a generally good correspondence between the predicted and actual tuning and demonstrate a relatively uniform distribution of tuning across the tested range of frequencies.

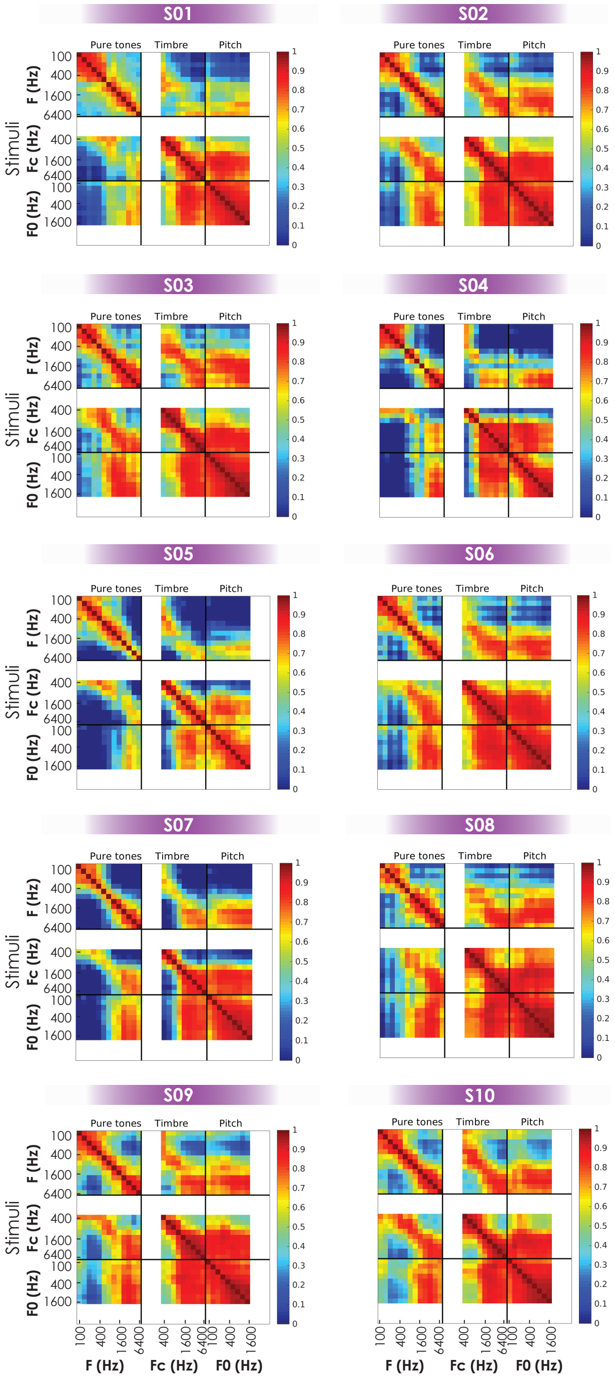


**Figure S5. Individual Representational Similarity Matrices (RSMs)** The RSMs within the ROI of each participant, averaged across repeats for a given stimulus, and thresholded to include only voxels with a GLM $R^2$ of at least 10%. White spaces indicate pure-tone frequencies that do not have corresponding Fc or F0 values for the timbre and pitch conditions, respectively.

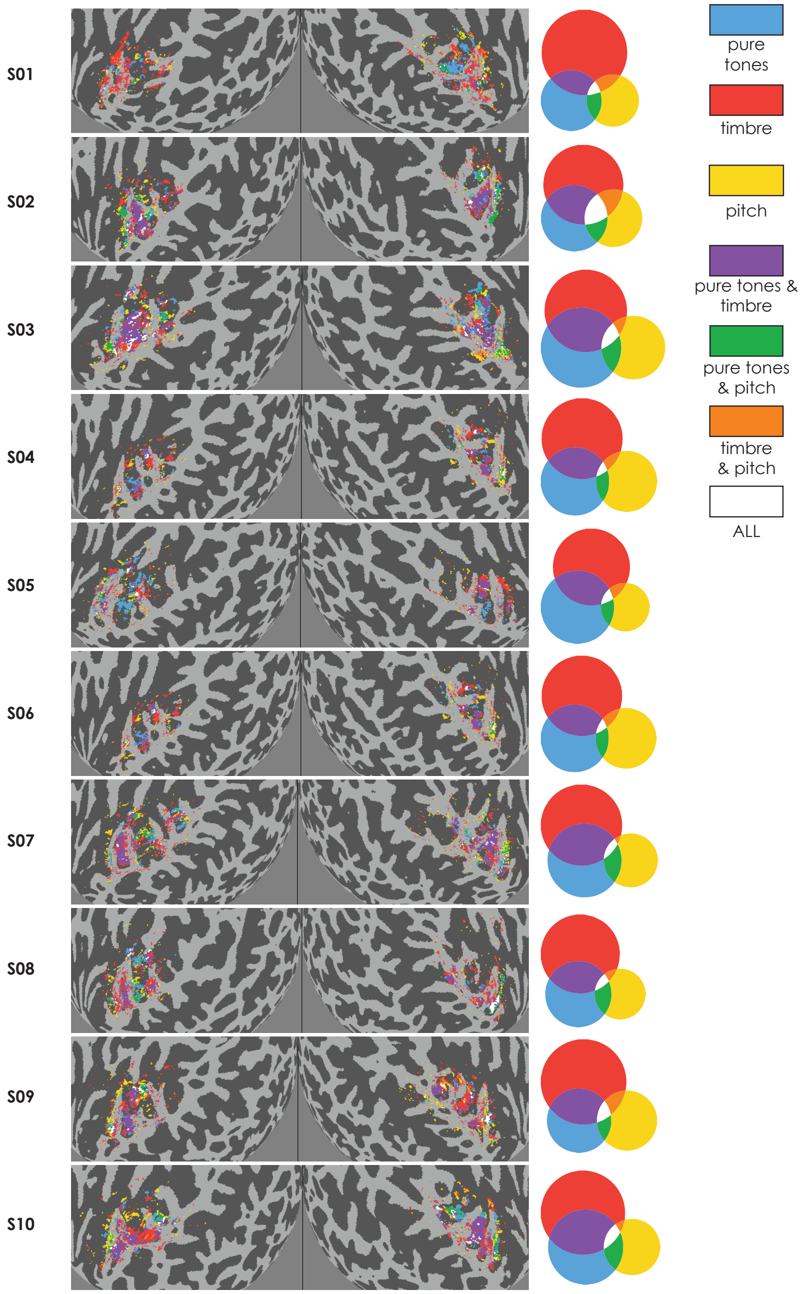


**Figure S6. Joint Tuning Surface Plots and Venn Diagrams for Each Participant**

Surface maps for each participant showing voxels with a model $R^2$ of at least 30% for one or more features within the feature tuning model. The adjacent Venn diagrams show the proportion of voxels exhibiting tuning sensitivity to one or more features, also only including voxels with a model $R^2$ of at least 30%. See legend for a description of the color map.

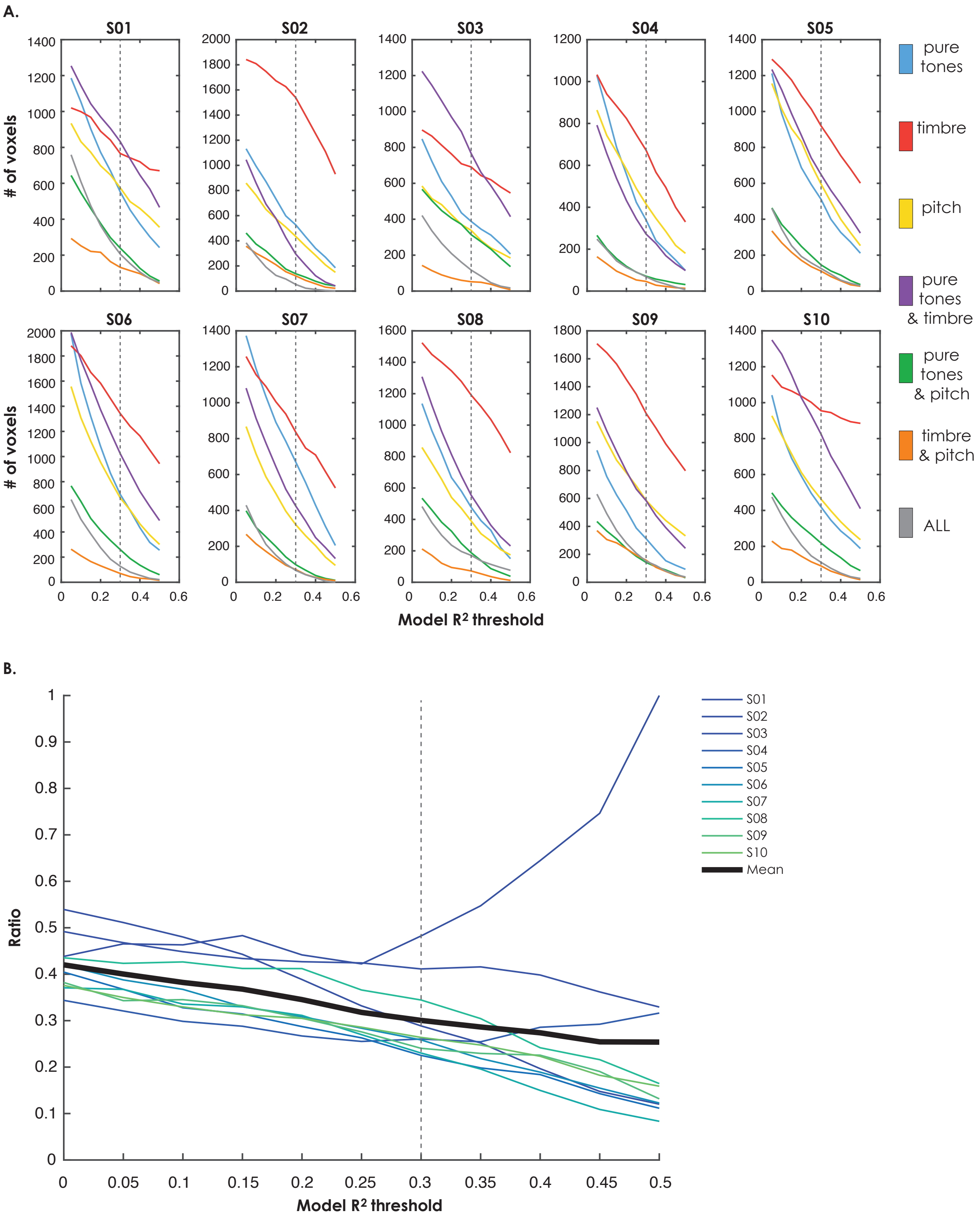


**Figure S7. Venn Diagram Characteristics as a Function of $R^2$ Threshold**

(A) Number of voxels in each section of the Venn diagram for each participant as a function of the feature tuning model $R^2$ threshold. Each panel represents a different participant. Different colors correspond to the different sections of the Venn diagrams, as shown in the legend. Dashed lines denote the threshold (30%) used for the Venn diagrams (Figures 7 and S6).

(B) Ratio of the pure-tone and pitch to pure-tone and timbre sections of the Venn diagrams as a function of the feature tuning model $R^2$ threshold. Each line represents data from a different participant. The black line shows the mean ratio across participants. Dashed line denotes the threshold used for the Venn diagrams (Figures 7 and S6).

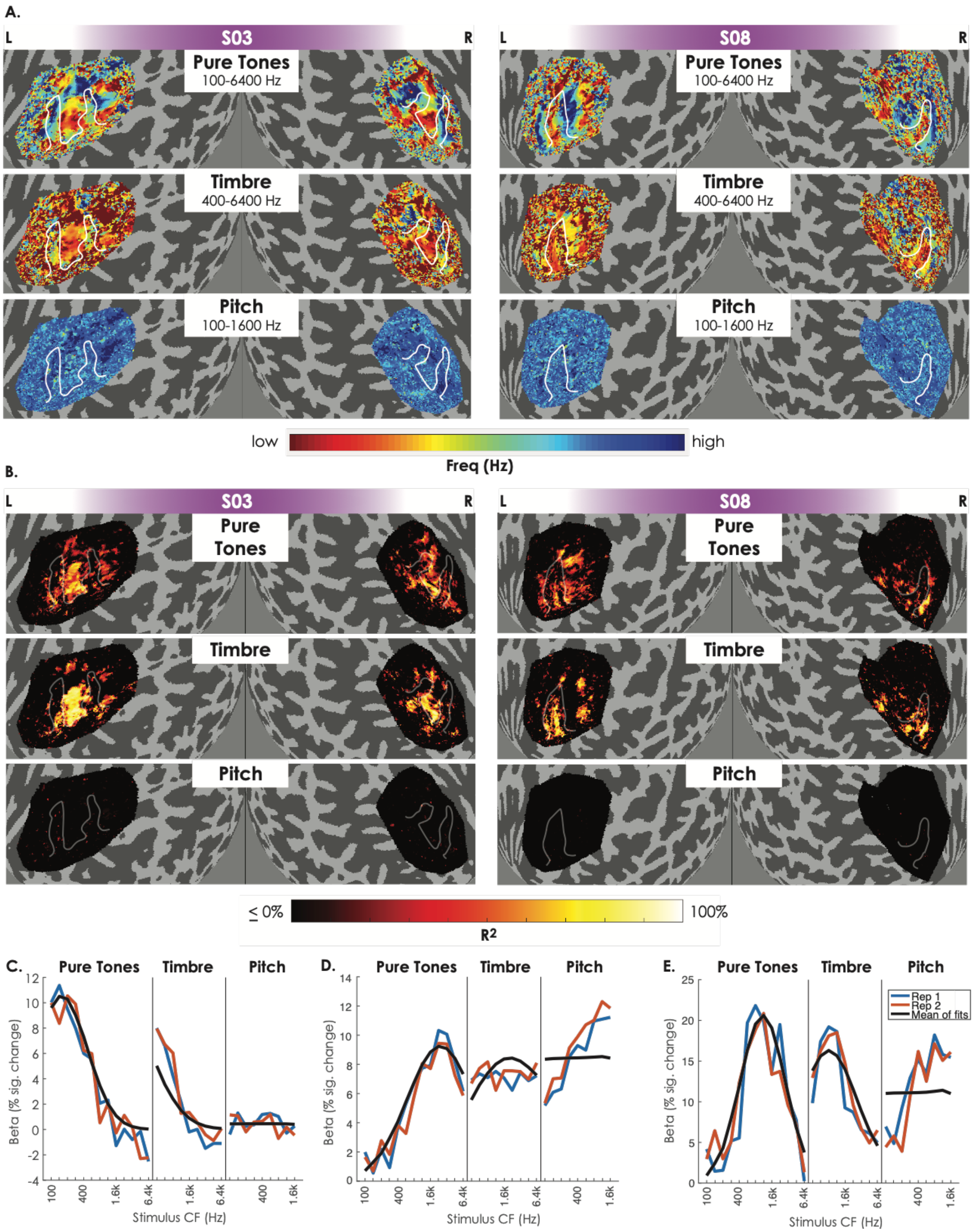


**Figure S8. The Spectral Tuning Model is a Poor Model of Pitch**

(A) Unthresholded CF maps for the spectral tuning model for two individual participants. Maps are shown on inflated cortical spheres of the left and right hemispheres, respectively.

Each row is the CF map for a given condition (pure tones, timbre, and pitch, respectively). Custom color maps span the respective CF range of each condition, as labeled. L = left hemisphere, R = right hemisphere. HG denoted by white lines.

(B) $R^2$ heat maps for the spectral tuning model for the same two participants as in Panel A.

(C) A voxel for a single participant showing good tuning for pure tones and timbre, but poor tuning for pitch. The black line is the model prediction. The colored lines are the beta values for each stimulus type, ordered from low to high.

(D) A voxel for a single participant showing good tuning for pure tones and pitch, but poor tuning for timbre. Same plotting conventions as in Panel C.

(E) A voxel for a single participant showing good tuning for all three conditions but demonstrating that the spectral tuning model is unable to account for the tuning in the pitch condition. Same plotting conventions as in Panels C and D.

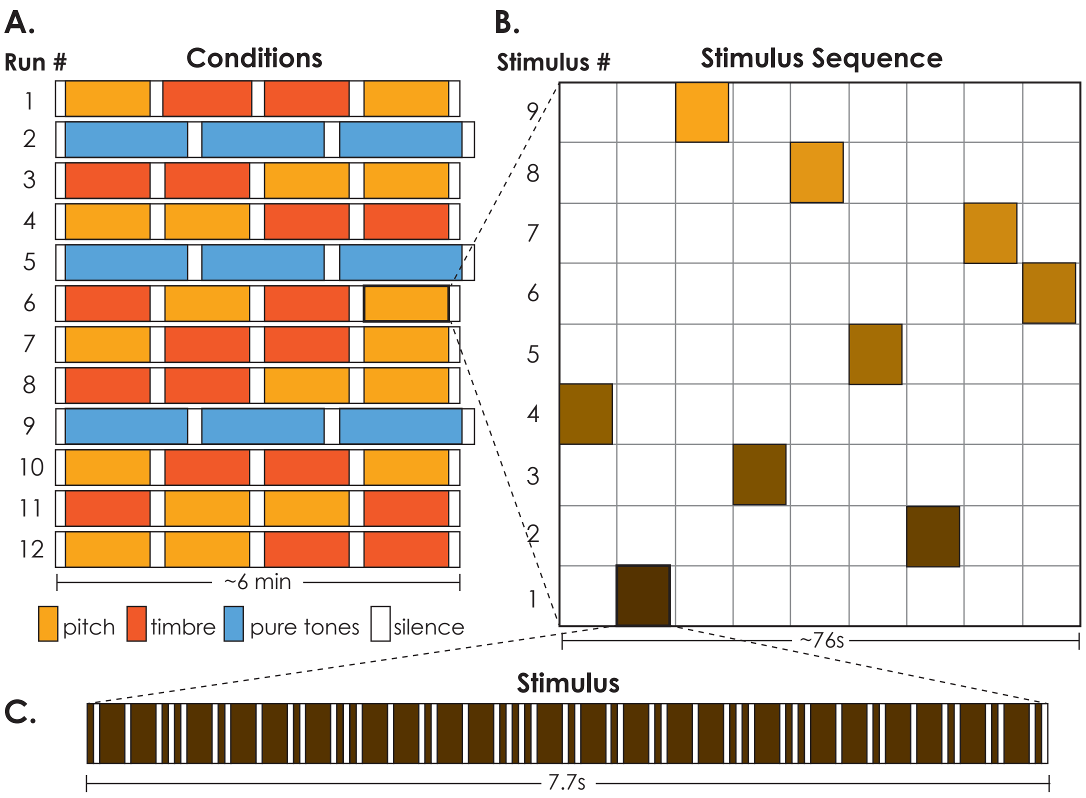


**Figure S9. Schematic of Functional Runs within a Scan Session**

(A) The order of the conditions within a run was pseudo-randomized and the order of runs across the session was counterbalanced across participants.

(B) Pseudo-randomized tone sequence steps within a condition block.

(C) Pseudo-randomized sequence of tones with two different durations to create "Morse-code"-like rhythm.
